# The SARS-CoV-2 mRNA-1273 vaccine elicits more RBD-focused neutralization, but with broader antibody binding within the RBD

**DOI:** 10.1101/2021.04.14.439844

**Authors:** Allison J. Greaney, Andrea N. Loes, Lauren E. Gentles, Katharine H.D. Crawford, Tyler N. Starr, Keara D. Malone, Helen Y. Chu, Jesse D. Bloom

**Affiliations:** Basic Sciences Division and Computational Biology Program, Fred Hutchinson Cancer Research Center; Seattle, WA 98109, USA; Department of Genome Sciences & Medical Scientist Training Program, University of Washington; Seattle, WA 98195, USA; Howard Hughes Medical Institute; Chevy Chase, MD 20815, USA; Department of Microbiology, University of Washington; Seattle, WA 98195, USA; Division of Allergy and Infectious Diseases, University of Washington; Seattle, Washington, USA

## Abstract

The emergence of SARS-CoV-2 variants with mutations in key antibody epitopes has raised concerns that antigenic evolution will erode immunity. The susceptibility of immunity to viral evolution is shaped in part by the breadth of epitopes targeted. Here we compare the specificity of antibodies elicited by the mRNA-1273 vaccine versus natural infection. The neutralizing activity of vaccine-elicited antibodies is even more focused on the spike receptor-binding domain (RBD) than for infection-elicited antibodies. However, within the RBD, binding of vaccine-elicited antibodies is more broadly distributed across epitopes than for infection-elicited antibodies. This greater binding breadth means single RBD mutations have less impact on neutralization by vaccine sera than convalescent sera. Therefore, antibody immunity acquired by different means may have differing susceptibility to erosion by viral evolution.

**One Sentence Summary:** Deep mutational scanning shows the mRNA-1273 RBD-binding antibody response is less affected by single viral mutations than the infection response.

## INTRODUCTION

Mitigation of the SARS-CoV-2 pandemic will depend on population immunity acquired via infection or vaccination. Unfortunately, humans are repeatedly re-infected with the endemic “common-cold” coronaviruses (*1*), at least in part because these viruses evolve to escape neutralizing antibody immunity elicited by prior infection (*2*). SARS-CoV-2 is already undergoing similar antigenic evolution, with the recent emergence of new viral lineages with reduced neutralization by antibodies elicited by infection and vaccination (*3*–*8*). Preliminary results suggest that immunity still provides substantial protection against infection and severe disease (*9*, *10*) caused by these new viral lineages—but if SARS-CoV-2 is similar to other human coronaviruses, then at minimum the protection against reinfection will eventually be eroded by viral evolution.

However, unlike for other human coronaviruses, a large fraction of the population is acquiring SARS-CoV-2 immunity from vaccination rather than infection. The first two vaccines approved for emergency use in the United States were Moderna’s mRNA-1273 and Pfizer/BioNTech’s BNT162b2. Both mRNA vaccines encode the full SARS-CoV-2 spike ectodomain with a transmembrane anchor and stabilizing S-2P mutations (*11*). It is possible that these vaccines could elicit antibodies with distinct specificities compared to natural infection due to variation in the spike (e.g., the S-2P mutations) or divergent immune responses to a two-dose mRNA vaccine versus infection. If the specificities differ, this could influence the impact of viral evolution on SARS-CoV-2 immunity.

Here we use a combination of serological assays and deep mutational scanning to map the specificity of the human polyclonal antibody response to the mRNA-1273 vaccine. The vaccine elicits neutralizing activity that is even more focused on the spike receptor-binding domain (RBD) than infection-elicited immunity. However, within the RBD, binding by vaccine-elicited antibodies is usually less affected by single mutations. As a result, common RBD mutations sometimes eliminate less of the neutralizing activity of mRNA-1273 vaccine sera than convalescent sera—and vaccine sera retain substantial RBD-directed neutralization even in the presence of mutations to three major RBD neutralizing epitopes. These findings suggest that antibody immunity acquired by infection and mRNA vaccines may have different sensitivity to erosion by viral evolution.

## RESULTS

### Sera from individuals vaccinated with the Moderna spike mRNA vaccine, mRNA-1273

We studied sera from adults (ages 18–55 years) who received two doses of the Moderna mRNA-1273 vaccine in the phase 1 clinical trials (*12*). The majority of our study focused on 14 individuals who received the 250 μg dose, although we validated key conclusions with a smaller subset of eight trial participants who received the 100 μg dose. The sera were collected at 36 and 119 days after the first vaccine dose, corresponding to 7 and 90 days after the second dose. It was previously shown that these individuals had high levels of binding and neutralizing antibodies against SARS-CoV-2, with neutralizing antibody titers within the upper quartile of sera from SARS-CoV-2 convalescent individuals (*12*).

Throughout, we compare key findings for vaccine sera to those for convalescent plasmas from two independent cohorts (*13*, *14*). The convalescent plasmas were characterized in earlier studies (*13*–*16*), and grouped into an early time point of 15–60 days post-symptom onset and a late time point of 100–150 days post-symptom onset.

### The neutralizing activity of mRNA-1273 vaccine-elicited antibodies is even more RBD-focused than for infection-elicited antibodies

The majority of the neutralizing activity of most convalescent sera and plasmas is due to RBD-binding antibodies (*15*, *17*, *18*). To determine if neutralization by vaccine sera is similarly RBD-targeted, we depleted RBD-binding antibodies from the day 36 and 119 sera from 14 individuals who received the 250 μg vaccine dose. We then measured serum IgG binding to the RBD and full spike ectodomain before and after depletion. As expected, depletion removed all RBD-binding antibodies (**Fig. 1A, S1A,B**). However, depleting RBD-binding antibodies only moderately decreased spike-binding activity (**Fig. 1B, S1B**), consistent with studies showing that a minority of spike-binding vaccine-elicited B cells target the RBD (*5*, *19*).

**Fig. 1.**
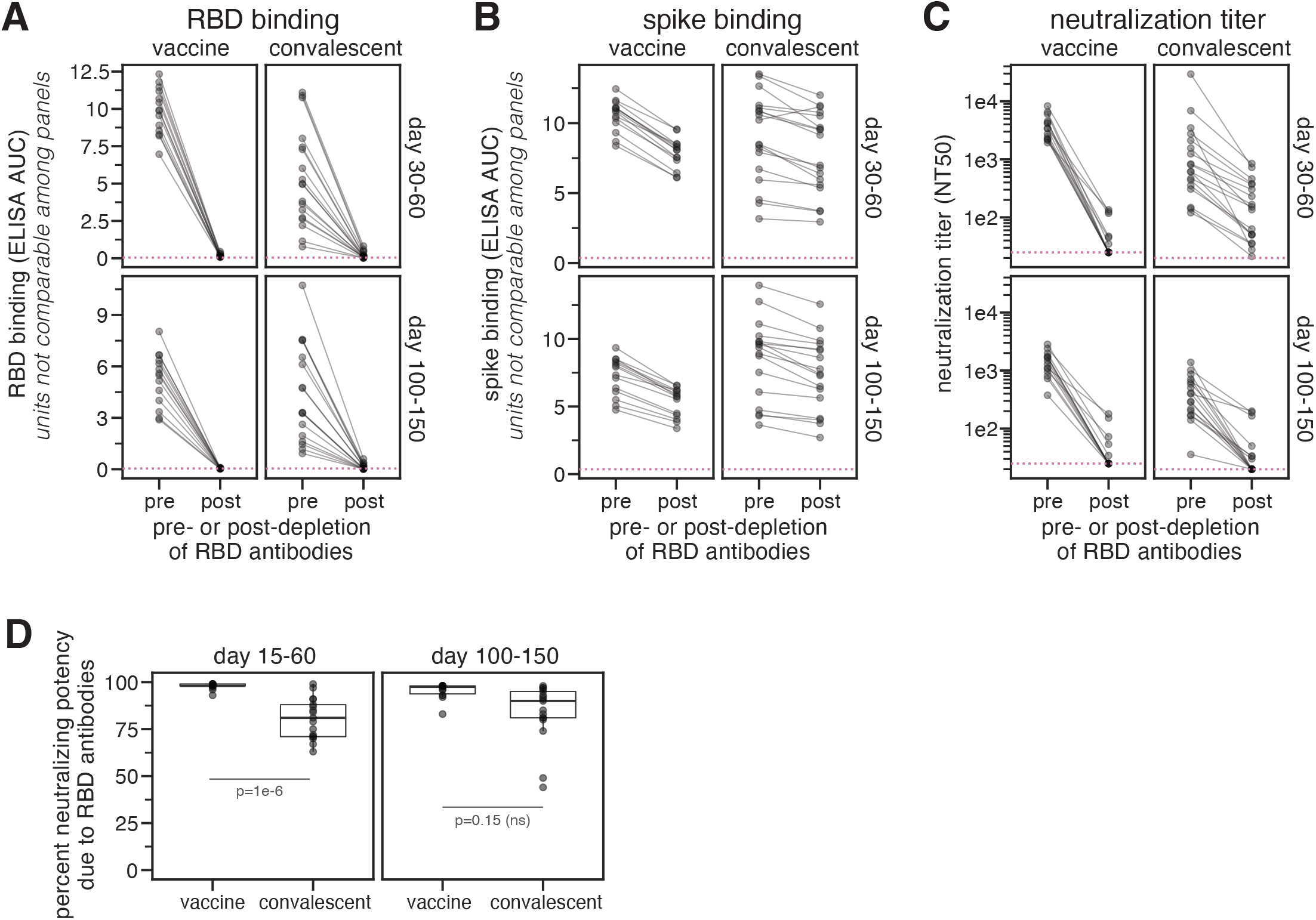
RBD-binding antibodies are responsible for most neutralizing activity of *mRNA-1273* vaccine-elicited sera. **(A)** Binding of serum antibodies to SARS-CoV-2 RBD, as measured by ELISA area-under-the-curve (AUC), for vaccine-elicited sera and convalescent plasmas before and after depletion of RBD-binding antibodies. The dashed pink line indicates binding of pre-pandemic sera. **(B)** Binding of serum antibodies to the full spike ectodomain. The y-axis scale units in (A) and (B) are not comparable between vaccine and convalescent samples due to different dilution factors (beginning at 1:500 for vaccine sera and 1:100 for convalescent plasmas). **(C)** Neutralization titer (reciprocal IC50) of vaccine-elicited sera and convalescent plasmas before and after depletion of RBD-binding antibodies. The limit of detection is shown as a dashed horizontal pink line. **(D)** Percent of neutralizing activity of vaccine-elicited sera and convalescent plasmas due to RBD-binding antibodies. P-values are from a log-rank test accounting for censoring. n=17 for each time point for convalescent plasmas, and n=14 for each time point for vaccine sera. All measurements of convalescent plasma binding and neutralization were previously reported in (*15*).

To determine the contribution of RBD-binding antibodies to neutralization, we measured the neutralizing activity of vaccine sera before and after depleting RBD-binding antibodies, using spike-pseudotyped lentiviral particles. For 13 of 14 vaccinated individuals, >90% of the neutralizing activity at both time points was due to RBD-binding antibodies (**Fig. 1C,D, Table S1**). For 17 of 28 vaccine sera, depletion of RBD-binding antibodies reduced the neutralization titer (reciprocal IC50) from >1000 to <25 (**Fig. 1C,D, S1C,D**). The percent neutralizing activity due to RBD-binding antibodies was higher for vaccine sera than for convalescent plasmas collected at similar time points (*15*) (**Fig. 1C,D**), although the difference was only statistically significant at the day 15–60 time point. These assays were performed in 293T cells over-expressing human ACE2, which may underestimate contributions of non-RBD-binding antibodies to viral neutralization (*6*, *20*, *21*). Nonetheless, because the same assay was used for vaccine and convalescent samples, we conclude that the neutralizing activity of the antibody response elicited by the mRNA-1273 vaccine is more focused on the RBD than for infection-elicited antibodies.

### Complete mapping of RBD mutations that reduce binding by vaccine-elicited sera at 119 days post-vaccination

We used deep mutational scanning (*15*, *22*) to map all mutations to yeast-displayed RBD that reduced vaccine serum antibody binding. Briefly, we incubated duplicate libraries of yeast expressing RBD mutants with each serum, and used fluorescence-activated cell sorting (FACS) to enrich cells expressing RBD mutants with reduced serum binding (**Fig. S2, S3, Table S2**). These libraries contained 3,804 of the 3,819 possible single amino-acid mutations to the Wuhan-Hu-1 RBD, 2,034 of which are tolerated for proper folding and at least modest ACE2 binding (*23*). We used deep sequencing to quantify the “escape fraction” for each of the 2,034 tolerated RBD mutations against each serum. These escape fractions range from 0 (no cells with the mutation in the serum-escape bin) to 1 (all cells with the mutation in the serum-escape bin). We represent the escape maps as logo plots, where the height of each letter is proportional to its escape fraction (**Fig. 2, Fig. S5, S6**).

**Fig. 2.**
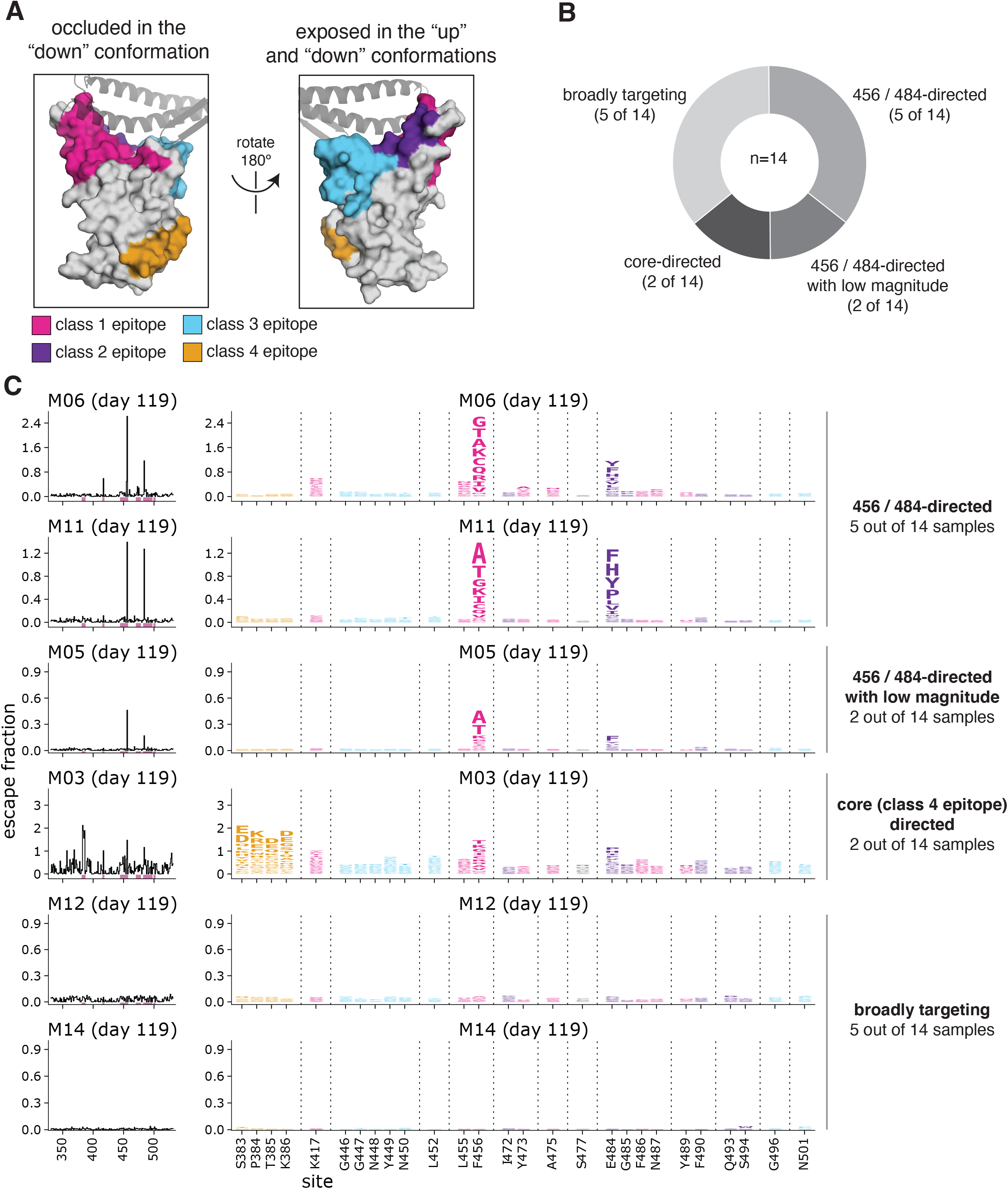
Complete maps of RBD mutations that reduce binding by serum collected 119 days post-vaccination with the 250 μg dose. **(A)** The epitopes of four major classes (*24*) of RBD-binding antibodies are colored on the RBD surface (PDB 6M0J). ACE2 is shown as a gray cartoon. **(B)** Number of sera that fell into each of the four major categories of binding-escape maps as categorized by subjective visual inspection. **(C)** Escape maps for six representative sera. The line plots at left indicate the sum of effects of all mutations at each RBD site on serum antibody binding, with larger values indicating more escape. The logo plots at right show key sites (highlighted in purple on the line plot x-axes). The height of each letter is that mutation’s escape fraction; larger letters indicate a greater reduction in binding. Escape fractions are not strictly comparable between samples due to the use of sample-specific FACS selection gates—therefore, for each sample, the y-axis is scaled independently (see **Methods**). RBD sites are colored by epitope as in (A). The escape fractions were well-correlated between independent libraries, and we report the average of duplicate measurements throughout (Table S3, Fig. S4). Escape maps for all 14 sera collected at day 119 from individuals who received the 250 μg dose are shown in **Fig. S5**. Comparable escape maps for eight individuals who received the 100 μg dose are in **Fig. S6**. Interactive versions of logo plots and structural visualizations are at https://jbloomlab.github.io/SARS-CoV-2-RBD_MAP_Moderna/.

The escape maps for sera collected at day 119 from individuals who received the 250 μg vaccine dose fell into four qualitative categories (**Fig. 2**). For 5 of 14 individuals, escape from antibody binding was focused on RBD sites 456 and 484 (**Fig. 2, S5**). These two sites are on the receptor-binding ridge in the “class 1” and “class 2” RBD epitopes, respectively (*24*) (**Fig. 2A**). Two more individuals also had escape maps that were focused on sites 456 and 484, but with a very low overall magnitude of escape (**Fig. 2, S5**). For 2 of 14 individuals, serum binding was most affected by mutations in the “class 4” epitope located in the core RBD, including sites 383 to 386 (**Fig. 2, S5**). The escape maps for the remaining 5 individuals were “flat,” meaning that no single mutation had a large effect on serum binding, suggestive of broad binding to multiple RBD epitopes (**Fig. 2, S5**).

To determine if the vaccine dose affected the RBD binding specificity of the polyclonal antibody response, we mapped binding escape from the day 119 sera from 8 individuals vaccinated with 100 μg rather than 250 ug doses. The escape maps of the 100 μg cohort resembled those of the 250 μg cohort and fell into the 456/484-targeting, core-targeting, or “flat” categories (**Fig. S6**). Although the sample sizes are small, and a higher fraction of the 100 μg dose escape maps were “flat” than for the 250 μg cohort (4/8 versus 5/14, respectively), this suggests 100 and 250 ug doses elicit antibody responses similar in the breadth of their RBD binding specificity.

### Binding escape maps become more focused from 36 days to 119 days post-vaccination

To examine longitudinal changes in binding specificity of vaccine-elicited serum antibodies to the RBD, we also determined binding-escape maps for sera collected at day 36 post-vaccination from five individuals who received the 250 μg dose (**Fig. 3**). All of these day-36 sera had relatively “flat” escape maps, meaning that no single mutation had a large effect on serum binding. But by day 119, the escape maps for most individuals were more focused on specific sites in the RBD. Specifically, for four of five individuals, the escape maps became focused on RBD sites 456 and 484 (**Fig. 3**). For one of these individuals, the focusing on sites 456 and 484 was accompanied by increased focusing on the class 4 epitope, including sites 383–386. Only one individual had a day-119 escape map as flat as the day-36 escape map. These results suggest that as the vaccine-induced RBD-binding antibody response matures over time, it becomes more focused on specific sites in the RBD.

**Fig. 3.**
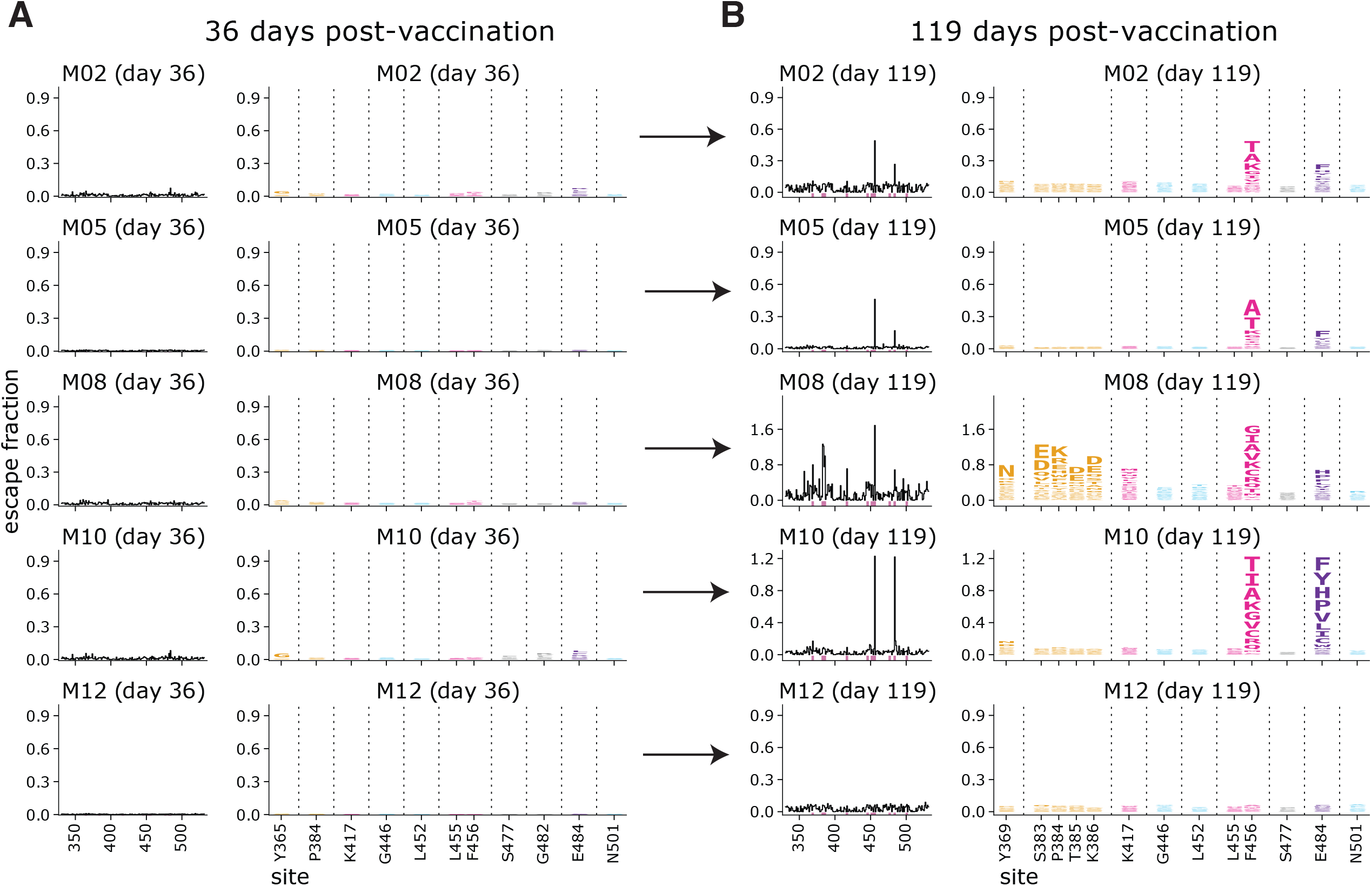
Comparison of escape maps for sera collected at day 36 and day 119 post-vaccination shows that the RBD-binding response focuses over time. Escape maps for sera at day 36 **(A)** and day 119 **(B)** from 5 individuals who received the 250 μg vaccine dose. The day 36 maps are all relatively flat, indicating no RBD mutation has a large effect on serum antibody binding. By day 119, the maps are often more focused on sites 456 and 484. The y-axis is scaled separately for each serum (see **Methods**). Interactive versions are at https://jbloomlab.github.io/SARS-CoV-2-RBD_MAP_Moderna/.

### RBD binding by vaccine-elicited sera is broader than for convalescent plasmas

To elucidate differences in the specificity of the RBD-binding antibody response elicited by vaccination versus infection, we compared the vaccine-sera escape maps to ones that we previously determined for convalescent plasmas (*15*, *16*). At both 15–60 and 100–150 day time points, the convalescent escape maps were more focused on specific RBD sites than the vaccine escape maps (**Fig. 4A**). The difference was especially striking at the early time point, where the day 36 vaccine samples all had flat escape maps, whereas the convalescent samples often had escape maps indicating that antibody binding was strongly affected by mutations at specific RBD sites such as 456 and 484 (**Fig. 4A**). The difference between the vaccine and convalescent samples was less striking at the later time point, but the convalescent maps were still more focused than the vaccine maps. There were also some differences in the RBD sites where mutations affected binding for the vaccine versus convalescent samples. While most samples of both types were affected by mutations at sites 456 and 484, the convalescent samples tended to also be affected by mutations to the 443–450 loop in the class 3 epitope, whereas mutations in the class 4 epitope spanning sites 383–386 sometimes had a more pronounced effect on the vaccine samples **(Fig. 4A, 2, S5**).

**Fig. 4.**
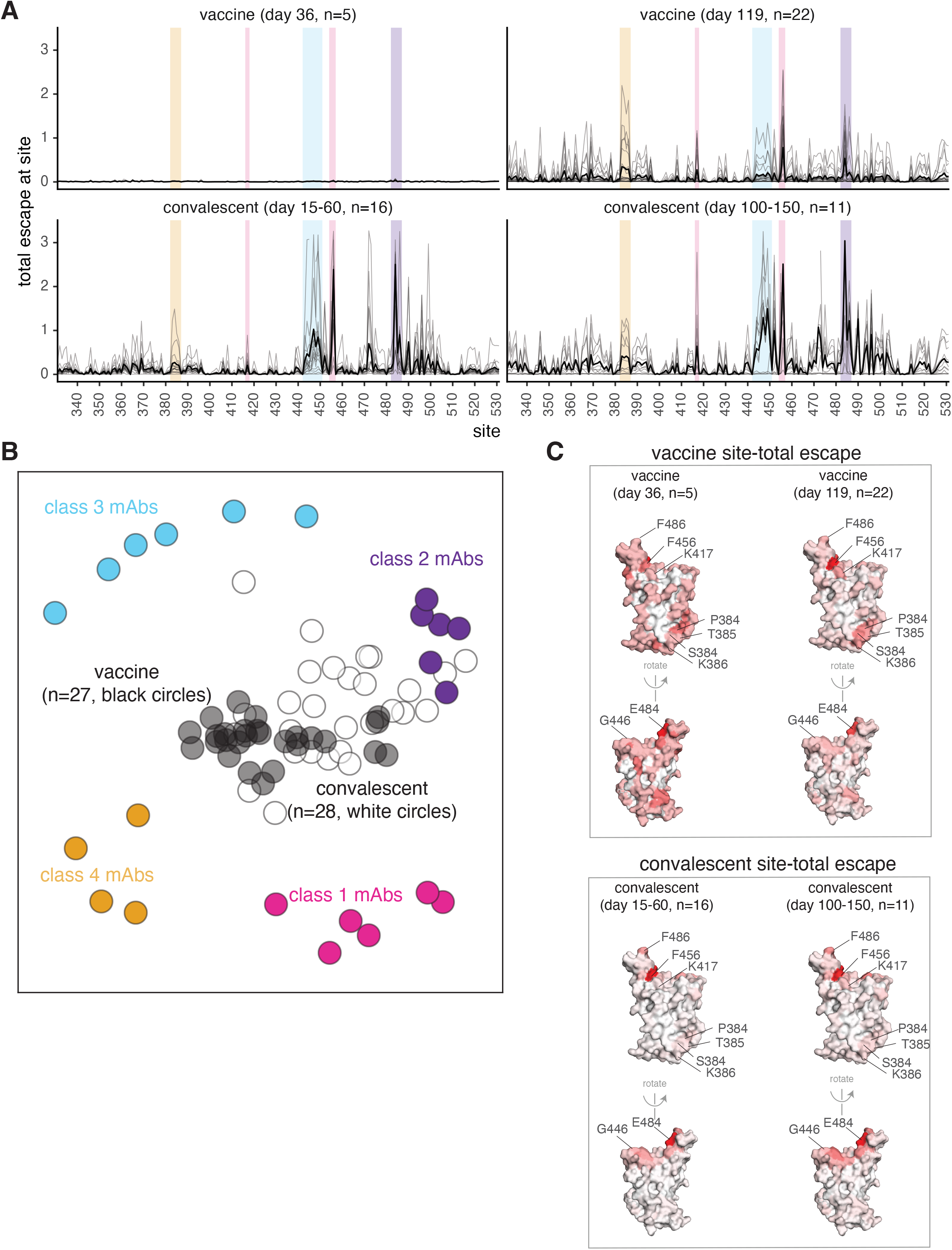
The binding of vaccine-elicited polyclonal antibodies is more broadly distributed across the RBD than for infection-elicited antibodies. **(A)** Escape from RBD-binding antibodies at each site in the RBD for vaccine sera or convalescent plasmas collected at early or late time points. Thin gray lines show individual serum or plasma samples, and the thick black line shows the mean (number of samples is indicated in the plot titles). Key sites within the epitopes of each major RBD antibody class are highlighted with the colors defined in **Fig. 2A** and in panel (B). **(B)** Relationships among escape maps of vaccine sera, convalescent plasmas, and monoclonal antibodies visualized with a multidimensional scaling projection. Vaccine sera include both doses and time points. Convalescent plasmas include all time points. **(C)** Binding escape at each site mapped onto the RBD surface after averaging across all sera/plasmas in each group. The RBD surface coloring is scaled from white to red, with white indicating no escape, and red indicating the site with the greatest escape. The color scaling spans the full range of white to red for each serum/plasma group, so the quantitative scale is not comparable across groups. Escape maps for monoclonal antibodies previously described in (*16*, *22*, *25*–*27*), and convalescent plasmas in (*15*, *16*).

To visualize relationships between vaccine- and infection-elicited antibody responses, we used multidimensional scaling to create a two-dimensional projection of the escape maps for the vaccine sera, convalescent plasmas (*15*, *16*), and previously characterized monoclonal antibodies (*16*, *22*, *25*–*27*) (**Fig. 4B**). In this projection, antibodies or sera/plasmas with similar binding-escape mutations are located close together, while those affected by distinct mutations are far apart. As previously reported (*16*), convalescent plasmas clustered closest to class 2 antibodies (**Fig. 4B**), which are generally most affected by mutations to site 484. The vaccine sera, on the other hand, were more centrally located in the middle of the antibodies of all four classes, reflecting their flatter binding-escape maps that were less dominated by mutations that escape any single antibody class (**Fig. 4B**).

To examine sites of binding-escape mutations in the context of the RBD’s structure, we projected the total escape at each site averaged across all vaccine or convalescent samples at each time point onto the surface of the RBD (**Fig. 4C**). The sites where mutations affected binding of vaccine sera were broadly distributed across the RBD surface (**Fig. 4C**), whereas convalescent plasmas were most affected by mutations at just a few key regions (sites 456 and 484, and to a lesser degree the 443–450 loop) (**Fig. 4C**). However, as noted above, binding escape from the vaccine sera was somewhat more focused at day 119 relative to day 36, including at sites 456, 484, and 383–386.

### Single RBD mutations have less impact on vaccine-elicited than infection-elicited neutralizing activity

We tested key RBD mutations in spike-pseudotyped lentiviral neutralization assays against a subset of vaccine and convalescent sera. We used the binding-escape maps to choose six representative samples each from the day 100–150 vaccine and convalescent sera for which >90% of the neutralizing activity was due to RBD-binding antibodies (**Fig. 1, S1**, (*15*)). The escape maps for the vaccine and convalescent samples chosen for these assays are summarized in **Fig. 5A** and detailed in **Fig. 2** and **Fig. S7**.

**Fig. 5.**
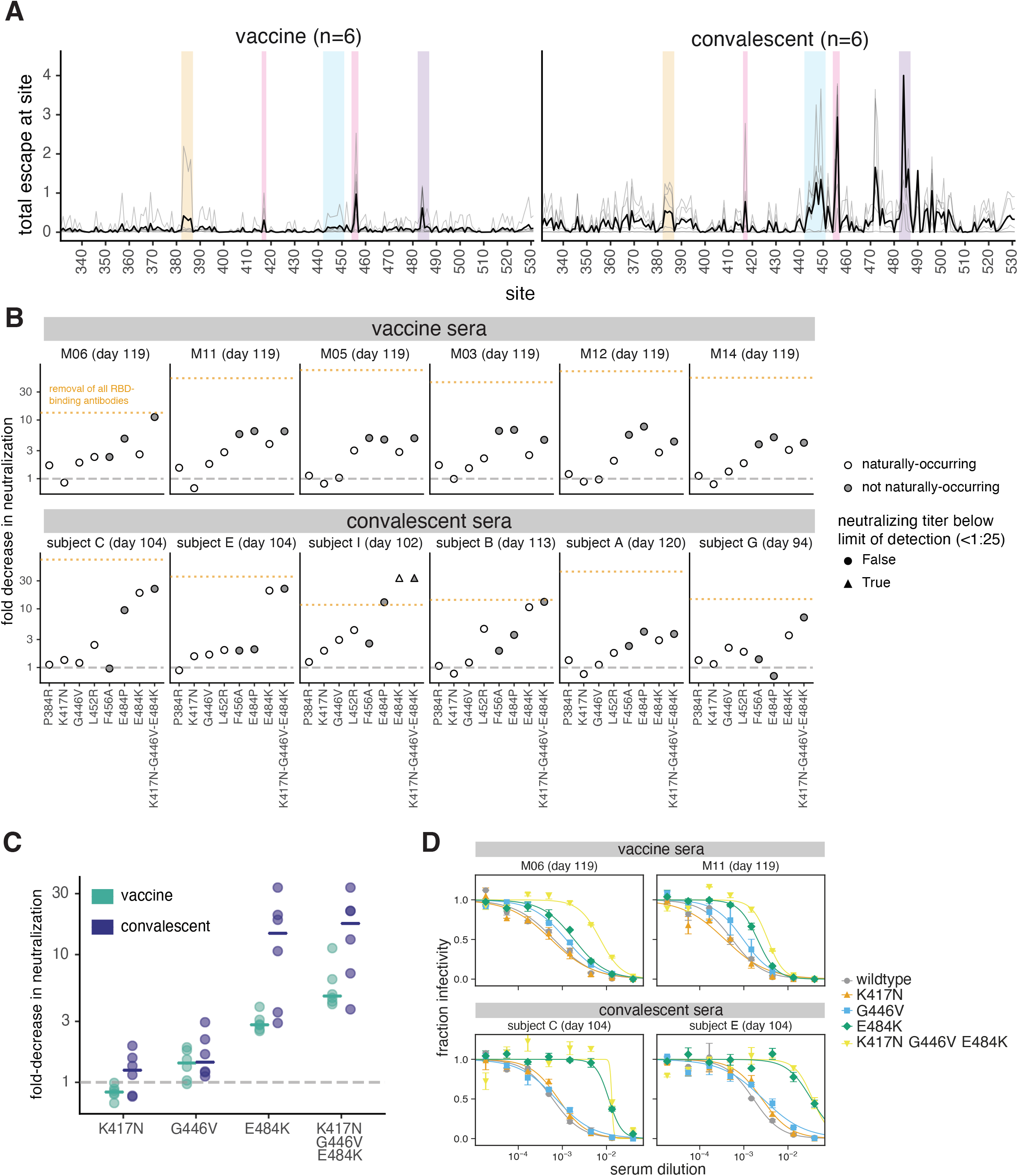
Effects of RBD mutations on neutralization by day 100–150 sera from vaccinated and convalescent individuals. **(A)** Total binding escape at each RBD site is shown for the six vaccine and convalescent samples tested in neutralization assays. The thin gray lines show individual samples, and the dark black line shows the mean. Key sites within each epitope are highlighted using the same color scheme as in **Fig. 2A**. **(B)** Neutralization of G614 spike-pseudotyped lentiviral particles with the indicated RBD mutations, shown as the fold-increase in IC50 compared to G614 spike with no additional mutations. Mutations that have been observed in human SARS-CoV-2 isolates are colored in white, and non-naturally-occurring mutations in gray. The orange dashed line represents the effect of depleting all RBD-binding antibodies, as in **Fig. 1**. **(C)** The fold decrease in neutralization titer caused by individual mutations in each of the three major neutralizing epitopes of the RBD (K417 in the class 1 epitope, E484K in the class 2 epitope, and G446V in the class 3 epitope), as well as the combination of all three mutations. Horizontal lines represent the median. In (B) and (C), the dashed gray line indicates no change in neutralization relative to unmutated spike. **(D)** Representative neutralization curves from two vaccine and two convalescent samples against the triple mutant and its composite single mutations. IC50s and pseudovirus entry titers are shown in **Fig. S8** and all neutralization curves are in **Fig. S9**.

We performed neutralization assays on mutants in each of the four major RBD epitopes (class 1, K417N and F456A; class 2, E484P and E484K; class 3, G446V and L452R; class 4, P384R). Among these mutations, K417N, L452R, and E484K are present in emerging viral lineages (*28*–*33*) that have been shown to have reduced neutralization (*3*, *5*–*8*, *34*, *35*). We also tested a triple mutant, K417N-G446V-E484K, with mutations in the class 1, 2, and 3 epitopes.

For many convalescent sera, single RBD mutations reduced neutralization by approximately the same amount as removing all RBD-binding antibodies (**Fig. 5B, S8, S9**). However, no single RBD mutation we tested had a comparably large effect on vaccine sera (**Fig. 5B**). This result is consistent with the binding-escape maps, which generally indicate that vaccine sera have a broader RBD-binding specificity than convalescent sera.

The mutations that most impacted neutralization also differed between vaccine and convalescent sera (**Fig. 5B**). For convalescent sera, the largest reduction in neutralization was consistently caused by mutations to site E484 in the class 2 epitope (*16*, *22*), including the E484K mutation present in multiple emerging viral lineages (*28*, *31*, *33*). But for vaccine sera, E484K generally caused a more moderate decrease in neutralization. For some vaccine sera, another mutation at site E484 (E484P) caused a larger loss of neutralization, but E484P is not found in any sequenced isolates and reduces both ACE2 binding affinity (*23*) and viral entry titers (**Fig. S8D**). The F456A mutation to the class 1 epitope often reduced neutralization by vaccine sera, although it had little effect on convalescent sera; this mutation is also not observed in natural sequences and reduces viral entry titers (**Fig. S8D**). Mutations to the class 3 epitope (G446V, L452R) modestly reduced neutralization by some vaccine and convalescent sera (**Fig. 5B**). However, P384R in the less-neutralizing core RBD class 4 epitope (*17*, *18*, *36*, *37*) and K417N in the class 1 epitope had little effect on neutralization by any sera, consistent with previous reports (*5*–*7*, *38*). Importantly, while single mutations sometimes caused large decreases in neutralization by convalescent sera, in no case did they reduce neutralization by vaccine sera >10-fold or to a titer <100 (**Fig. 5B, S8**).

The fact that single mutations ablate the anti-RBD neutralizing activity of some convalescent sera but only modestly erode the activity of vaccine sera suggests that the vaccine elicits neutralizing antibodies with a greater number of RBD specificities. To test this idea, we performed neutralization assays with a triple mutant (K417N-G446V-E484K) containing a mutation in each of the class 1, 2, and 3 epitopes. For convalescent sera, the E484K mutation alone often caused a decrease in neutralization comparable to the triple mutant (**Fig. 5C,D, S8**), consistent with the convalescent escape maps showing a strong focus on site E484. In contrast, for vaccine sera, the triple mutant always reduced neutralization more than any of its constituent single mutations (**Fig. 5C,D, S8**). Moreover, for only one of the six vaccinated individuals did the triple mutant decrease neutralization to the same extent as removing all RBD-binding antibodies (**Fig. 5B**), indicating that the vaccine usually induces some neutralizing antibodies not escaped by mutations to sites K417, G446, and E484. These results are consistent with the escape maps indicating that the vaccine sera often have a broader RBD-binding specificity. Note that in some cases infection also elicited very broad anti-RBD neutralizing activity: for instance, serum from the convalescent individual with the broadest escape map (subject G, day 94) was substantially more affected by the triple mutant than any of its constituent single mutants (**Fig. 5B, S7, S8**).

## DISCUSSION

We have shown clear differences in the specificity of polyclonal serum antibodies acquired by infection versus vaccination with Moderna mRNA-1273. The neutralizing activity of vaccine sera is even more focused on the RBD than for convalescent sera, with the majority of vaccine sera losing all detectable neutralization at a 1:25 cutoff after depletion of RBD-directed antibodies. This fact is surprising, since the mRNA-1273 vaccine encodes the full spike ectodomain (*11*), and one conjectured benefit of full-spike versus RBD-only vaccines was elicitation of neutralizing antibodies targeting non-RBD subdomains.

At first glance, the RBD focusing of the vaccine sera neutralization might seem like a “bad” thing, but the rest of our results suggest that this may not be the case. Our comprehensive maps of how RBD mutations reduce serum antibody binding show that vaccine-induced antibodies are usually less affected than infection-elicited antibodies by any single RBD mutation. While infection-elicited RBD antibodies are often strongly focused on an epitope including site E484, vaccine-elicited antibodies bind more broadly across the RBD including to the more conserved “core”. This broader binding makes neutralization by vaccine sera more resistant to RBD mutations. For instance, RBD-directed neutralization by convalescent sera was greatly reduced or even eliminated by a combination of key mutations at the three major epitopes in the RBD’s receptor-binding motif, but all vaccine sera that we tested retained substantial neutralization against this triple mutant. This result implies that either vaccination induces an antibody response more broadly distributed across the RBD surface, or that the individual antibodies elicited by vaccination are more robust to these mutations (*39*, *40*).

Our results do not explain why the vaccine neutralizing response is so RBD-directed, but we note two possibilities. First, the vaccine encodes a stabilized S-2P spike, which could present some epitopes in slightly different conformations. Second, the vaccine is delivered by an mRNA-lipid nanoparticle, which may lead to different kinetics of antigen presentation than viral infection (*41*, *42*). Indeed, another recent study suggests that mRNA vaccination elicits a different distribution of isotypes and fewer antibodies that cross-react to common-cold coronaviruses compared to infection (*43*). A caveat is that some differences could also be because the vaccinated individuals were relatively young (18–55 years) and healthy, whereas the convalescent individuals were older (23–76 years, median 56) with a range of comorbidities (*13*).

More generally, our findings suggest that it is important to differentiate antibody immunity acquired by different means when assessing the impact of viral evolution. Significant effort is being expended to identify emerging antigenic variants of SARS-CoV-2 and determine which ones might evade immunity (*3*, *7*, *8*, *33*). Our findings suggest that the results could vary depending on the source of immunity. Furthermore, carefully characterizing and comparing the specificity of antibody immunity elicited by additional vaccine modalities could provide a basis for determining whether some will be more resistant to viral evolution.

## MATERIALS AND METHODS

### Study design and SARS-CoV-2 vaccine sera and convalescent plasmas

De-identified post-vaccination sera were obtained as secondary research samples from the National Institutes of Allergy and Infectious Diseases-sponsored mRNA-1273 phase 1 clinical trial (NCT04283461) (*12*). We obtained samples from 14 individuals who received two 250 μg doses of the mRNA-1273 vaccine, and 8 individuals who received two 100 μg doses. All individuals were between ages 18 and 55 years old. The samples were collected under the human subject approvals described in (*12*). Due to the de-identified nature of the samples, the work described in this paper was deemed non-human subjects research by the Fred Hutchinson Cancer Research Center Institutional Review Board.

Previously reported results from samples from two cohorts of SARS-CoV-2 convalescent individuals are reanalyzed here (*15*, *16*). One cohort of convalescent plasma samples were previously described (*13*, *15*) and collected as part of a prospective longitudinal cohort study of individuals with SARS-CoV-2 infection in Seattle, WA February–July 2020. The plasmas from 17 individuals were examined here (8/17 female; age range 23–76 years, mean 51.6 years, median 56 years). All data from this cohort (i.e., the neutralization and RBD- and spike-binding activity of plasmas pre- and post-depletion of RBD-binding antibodies in **Fig. 1** and RBD-binding escape maps in **Fig. 4**, **S6B**, and **S7**) were previously reported (*15*) with the exception of neutralization assays in **Fig. 5**, **S8**, and **S9**, which were newly performed in this study. This work was approved by the University of Washington Institutional Review Board.

All data from the second cohort of plasma samples (n=5) (i.e., the aggregated escape maps in **Fig. 4**) were previously reported (*16*) and are simply reanalyzed here. The plasmas were originally collected 21–35 days post-symptom onset as part of a prospective longitudinal cohort study of SARS-CoV-2 convalescent individuals in New York, NY, under the human subject approvals described in (*14*).

### Cell lines

HEK-293T (ATCC CRL-3216) cells were used to generate SARS-CoV-2 spike-pseudotyped lentiviral particles and 293T-ACE2 cells (BEI NR-52511) were used to titer the SARS-CoV-2 spike-pseudotyped lentiviral particles and to perform neutralization assays (see below).

### RBD deep mutational scanning library

The yeast-display RBD mutant libraries are previously described (*22*, *23*). Briefly, duplicate mutant libraries were constructed in the spike receptor binding domain (RBD) from SARS-CoV-2 (isolate Wuhan-Hu-1, Genbank accession number MN908947, residues N331-T531) and contain 3,804 of the 3,819 possible amino-acid mutations, with >95% present as single mutants. Each RBD variant was linked to a unique 16-nucleotide barcode sequence to facilitate downstream sequencing. As previously described, libraries were sorted for RBD expression and ACE2 binding to eliminate RBD variants that are completely misfolded or non-functional (i.e., lacking modest ACE2 binding affinity) (*22*).

### FACS sorting of yeast libraries to select mutants with reduced binding by polyclonal post-vaccination sera

Serum mapping experiments were performed in biological duplicate using the independent mutant RBD libraries, similarly to as previously described for monoclonal antibodies (*22*) and exactly as previously described for polyclonal plasmas (*44*). Briefly, mutant yeast libraries induced to express RBD were washed and incubated with serum at a range of dilutions for 1 h at room temperature with gentle agitation. For each serum, we chose a sub-saturating dilution such that the amount of fluorescent signal due to serum antibody binding to RBD was approximately equal across samples. The exact dilution used for each serum is given in **Supplementary Table 2**. After the serum incubations, the libraries were secondarily labeled with 1:100 FITC-conjugated anti-MYC antibody (Immunology Consultants Lab, CYMC-45F) to label for RBD expression and 1:200 Alexa-647-conjugated goat anti-human-IgA+IgG+IgM (Jackson ImmunoResearch 109-605-064) to label for bound serum antibodies. A flow cytometric selection gate was drawn to capture 3–6% of the RBD mutants with the lowest amount of serum binding for their degree of RBD expression (**Fig. S2, S3**). We also measured what fraction of cells expressing unmutated RBD fell into this gate when stained with 1× and 0.1× the concentration of serum. For each sample, approximately 10 million RBD+ cells (range 7.3e6 to 1.4e7 cells) were processed on the cytometer, with between 3e5 and 6e5 plasma-escaped cells collected per sample (see percentages in **Table S2**). Antibody-escaped cells were grown overnight in SD-CAA (6.7g/L Yeast Nitrogen Base, 5.0g/L Casamino acids, 1.065 g/L MES acid, and 2% w/v dextrose) to expand cells prior to plasmid extraction.

### DNA extraction and Illumina sequencing

Plasmid samples were prepared from 30 OD units (1.6e8 cfus) of pre-selection yeast populations and approximately 5 OD units (~3.2e7 cfus) of overnight cultures of serum-escaped cells (Zymoprep Yeast Plasmid Miniprep II) as previously described (*22*). The 16-nucleotide barcode sequences identifying each RBD variant were amplified by PCR and prepared for Illumina sequencing as described in (*23*). Barcodes were sequenced on an Illumina HiSeq 2500 with 50 bp single-end reads. To minimize noise from inadequate sequencing coverage, we ensured that each antibody-escape sample had at least 2.5× as many post-filtering sequencing counts as FACS-selected cells, and reference populations had at least 2.5e7 post-filtering sequencing counts.

### Analysis of deep sequencing data to compute each mutation’s escape fraction

Escape fractions were computed as described in (*22*), with minor modifications as noted below. We used the dms_variants package (https://jbloomlab.github.io/dms_variants/, version 0.8.5) to process Illumina sequences into counts of each barcoded RBD variant in each pre-sort and antibody-escape population using the barcode/RBD look-up table from (*23*).

For each serum selection, we computed the “escape fraction” for each barcoded variant using the deep sequencing counts for each variant in the original and serum-escape populations and the total fraction of the library that escaped antibody binding via the formula provided in (*22*). These escape fractions represent the estimated fraction of cells expressing that specific variant that falls in the escape bin, such that a value of 0 means the variant is always bound by serum and a value of 1 means that it always escapes serum binding. We then applied a computational filter to remove variants with low sequencing counts or highly deleterious mutations that might cause antibody escape simply by leading to poor expression of properly folded RBD on the yeast cell surface (*22*, *23*). Specifically, we removed variants that had (or contained mutations with) ACE2 binding scores < −2.35 or expression scores < −1, using the variant- and mutation-level deep mutational scanning scores from (*23*). Note that these filtering criteria are slightly more stringent than those used in (*22*) but are identical to those used in (*15*, *16*, *25*).

We next deconvolved variant-level escape scores into escape fraction estimates for single mutations using global epistasis models (*45*) implemented in the dms_variants package, as detailed at (https://jbloomlab.github.io/dms_variants/dms_variants.globalepistasis.html) and described in (*22*). The reported scores throughout the paper are the average across the libraries; these scores are also in **Supplementary Table 3**. Correlations in final single-mutant escape scores are shown in **Fig. S4**.

For plotting and analyses that required identifying RBD sites of “strong escape” (e.g., choosing which sites to show in logo plots in **Fig. 2, 3, S5, S6, S7**), we considered a site to mediate strong escape if the total escape (sum of mutation-level escape fractions) for that site exceeded the median across sites by >5-fold, and was at least 5% of the maximum for any site. Full documentation of the computational analysis is at https://github.com/jbloomlab/SARS-CoV-2-RBD_MAP_Moderna.

### Generation of pseudotyped lentiviral particles

We used spike-pseudotyped lentiviral particles that were generated essentially as described in (*46*), using a codon-optimized SARS-CoV-2 spike from Wuhan-Hu-1 that contains a 21-amino-acid deletion at the end of the cytoplasmic tail (*13*) and the D614G mutation that is now predominant in human SARS-CoV-2 (*47*). The plasmid encoding this spike, HDM_Spikedelta21_D614G, is available from Addgene (#158762) and BEI (NR-53765), and the full sequence is at (https://www.addgene.org/158762). Point mutations were introduced into the RBD of this plasmid via site-directed mutagenesis. Therefore, all mutations tested in this paper are in the G614 background, and are compared to a “wildtype” spike with G614.

To generate these spike-pseudotyped lentiviral particles (*46*), 6e5 HEK-293T (ATCC CRL-3216) cells per well were seeded in 6-well plates in 2 mL D10 growth media (DMEM with 10% heat-inactivated FBS, 2 mM l-glutamine, 100 U/mL penicillin, and 100 μg/mL streptomycin). 24h later, cells were transfected using BioT transfection reagent (Bioland Scientific, Paramount, CA, USA) with a Luciferase_IRES_ZsGreen backbone, Gag/Pol lentiviral helper plasmid (BEI NR-52517), and wildtype or mutant SARS-CoV-2 spike plasmids. Media was changed to fresh D10 at 24 h post-transfection. At ~60 h post-transfection, viral supernatants were collected, filtered through a 0.45 μm SFCA low protein-binding filter, and stored at −80°C.

### Titering of pseudotyped lentiviral particles

Titers of spike-pseudotyped lentiviral particles were determined as described in (*46*) with the following modifications. 100 μL of diluted spike-pseudotyped lentiviral particles was added to 1.25e4 293T-ACE2 cells (BEI NR-52511), grown overnight in 50 μL of D10 growth media in a 96-well black-walled poly-L-lysine coated plate (Greiner Bio-One, 655936). Relative luciferase units (RLU) were measured 65 h post-infection (Promega Bright-Glo, E2620) in the infection plates with a black back-sticker (Fisher Scientific, NC9425162) added to minimize background. Titers were first estimated from the average of 8, 2-fold serial dilutions of virus starting at 25 μL virus in a total volume of 150 μL, performed in duplicate, and normalized to a wild type D614G variant harvested on the same day. Quantitative titering was then performed at a single virus dilution, targeting 200,000 RLU per well. Values in Fig. S8D are an average RLUs per μL measured across 16 technical replicates at a single dilution.

### Neutralization assays

293T-ACE2 cells (BEI NR-52511) were seeded at 1.25e4 cells per well in 50 μL D10 in poly-L-lysine coated, black-walled, 96-well plates (Greiner 655930). 24 h later, pseudotyped lentivirus supernatants were diluted to ~200,000 RLU per well (determined by titering as described above and shown in **Fig. S5D** and incubated with a range of dilutions of serum for 1 h at 37 °C. 100 μL of the virus-antibody mixture was then added to cells. At ~70 h post-infection, luciferase activity was measured using the Bright-Glo Luciferase Assay System (Promega, E2610). Fraction infectivity of each serum antibody-containing well was calculated relative to a “no-serum” well inoculated with the same initial viral supernatant (containing wildtype or mutant RBD) in the same row of the plate. We used the neutcurve package (https://jbloomlab.github.io/neutcurve version 0.5.2) to calculate the inhibitory concentration 50% (IC50) and the neutralization titer 50% (NT50), which is simply 1/IC50, of each serum against each virus by fitting a Hill curve with the bottom fixed at 0 and the top fixed at 1. The full neutralization curves are in **Fig. S9**.

### Depletion of RBD-binding antibodies from polyclonal sera

Two rounds of sequential depletion of RBD-binding antibodies were performed for vaccine-elicited sera. Magnetic beads conjugated to the SARS-CoV-2 RBD (AcroBiosystems, MBS-K002) were prepared according to the manufacturer’s protocol. Beads were resuspended in ultrapure water at 1 mg beads/mL and a magnet was used to wash the beads 3 times in PBS with 0.05% BSA. Beads were then resuspended in PBS with 0.05% BSA at 1 mg beads per mL. Beads (manufacturer-reported binding capacity of 10–40 μg/mL anti-RBD antibodies) were incubated with human sera at a 3:1 ratio beads:serum (150 μL beads + 50 μL serum), rotating overnight at 4°C. A magnet (MagnaRack™ Magnetic Separation Rack, ThermoFisher CS15000) was used to separate antibodies that bind RBD from the supernatant, and the supernatant (the post-RBD antibody depletion sample) was removed. A mock depletion (pre-depletion sample) was performed by adding 150 μL of PBS + 0.05% BSA and incubating rotating overnight at 4°C. A second round of depletion was then performed to ensure full depletion of RBD-binding antibodies. For the neutralization assays on these sera depleted of RBD-binding antibodies shown in **Fig. S1D**; the reported serum dilution is corrected for the dilution incurred by the depletion process.

### Measurement of serum binding to RBD or spike by ELISA

The IgG ELISAs for spike protein and RBD were conducted as previously described (*48*). Briefly, ELISA plates were coated with recombinant spike and RBD antigens described in (*48*) at 2 μg/mL. Five 3-fold serial dilutions of sera beginning at 1:500 were performed in phosphate-buffered saline with 0.1% Tween with 1% Carnation nonfat dry milk. Dilution series of the “synthetic” sera comprised of the anti-RBD antibody rREGN10987 (*49*) or anti-NTD antibody r4A8 (*21*) and pooled pre-pandemic human serum from 2017-2018 (Gemini Biosciences; nos. 100–110, lot H86W03J; pooled from 75 donors) were performed such that the anti-spike antibody was present at a highest concentration of 0.25 μg/mL. Both antibodies were recombinantly produced by Genscript. The rREGN10987 is that used in (*25*) and the variable domain heavy and light chain sequences for r4A8 were obtained from Genbank GI 1864383732 and 1864383733 (*21*) and produced on a human IgG1 and IgK background, respectively. Pre-pandemic serum alone, without anti-RBD antibody depletion, was used as a negative control, averaged over 2 replicates. Secondary labeling was performed with goat anti-human IgG-Fc horseradish peroxidase (HRP) (1:3000, Bethyl Labs, A80-104P). Antibody binding was detected with TMB/E HRP substrate (Millipore Sigma, ES001) and 1 N HCl was used to stop the reaction. OD450 was read on a Tecan infinite M1000Pro plate reader. The area under the curve (AUC) was calculated as the area under the titration curve with the serial dilutions on a log-scale.

### Data visualization

The static logo plot visualizations of the escape maps in the paper figures were created using the dmslogo package (https://jbloomlab.github.io/dmslogo, version 0.6.2) and in all cases the height of each letter indicates the escape fraction for that amino-acid mutation calculated as described above. For each sample, the y-axis is scaled to be the greatest of (a) the maximum site-wise escape metric observed for that sample, (b) 20x the median site-wise escape fraction observed across all sites for that serum, or (c) an absolute value of 1.0 (to appropriately scale samples that are not “noisy” but for which no mutation has a strong effect on antibody binding). Sites K417, L452, S477, E484, and N501 have been added to logo plots due to their frequencies among circulating viruses. The code that generates these logo plot visualizations is available at https://github.com/jbloomlab/SARS-CoV-2-RBD_MAP_Moderna/blob/main/results/summary/escape_profiles.md.

In many of the visualizations (e.g., **Fig. 2, 3, 4, S5, S6, S7**), the RBD sites are categorized by epitope region (*24*) and colored accordingly. We define the class 1 epitope as residues 403+405+406+417+420+421+453+455–460+473–476+486+487+489+504, the class 2 epitope as residues 472+483–485+490–494, the class 3 epitope to be residues 345+346+437-452+496+498–501, and the class 4 epitope as residues 365–372+382–386.

For the static structural visualizations in the paper figures, the RBD surface (PDB 6M0J, (*50*)) was colored by the site-wise escape metric at each site, with white indicating no escape and red scaled to be the same maximum used to scale the y-axis in the logo plot escape maps, determined as described above. We created interactive structure-based visualizations of the escape maps using dms-view (*51*) that are available at https://jbloomlab.github.io/SARS-CoV-2-RBD_MAP_Moderna/. The logo plots in these escape maps can be colored according to the deep mutational scanning measurements of how mutations affect ACE2 binding or RBD expression as described above.

For the composite line plots shown in Fig. 4, the convalescent (day 15–60) group includes two independent cohorts of individuals, one recruited in New York, NY (n=5) (*14*), and another recruited in Seattle, WA (n=11) (*13*). The convalescent (day 100–150) group is from the longitudinal cohort recruited in Seattle, WA (n=11). The escape maps for convalescent individuals were previously reported in (*15*, *16*). The mRNA-1273 (day 119) group includes individuals who were vaccinated with either the 100 or 250 μg vaccine dose (n=8 and n=14, respectively). The y-axis maximum is scaled to 1.1 times the maximum group mean site-total escape among all groups, so outlier points exceeding this value are not shown.

### Quantification and Statistical Methods

In **Fig. 1D**, the percent of neutralizing activity of vaccine-elicited sera and convalescent plasma due to RBD-binding antibodies is shown as a Tukey boxplot (middle line=median, box limits=interquartile range) with individual measurements overlaid as points. P-values are from a log-rank test accounting for censoring, p=1.0 × 10^−6^ for convalescent vs. vaccine (day 15–60), and p=0.15 for convalescent vs. vaccine (day 100–150).

## Supplementary Materials

Fig. S1. Raw ELISA and neutralization curves of mRNA-1273 serum samples before and after depletion of RBD-binding antibodies.

Fig. S2. Schematic of the deep mutational scanning approach used to quantify the effects of RBD mutations on antibody escape.

Fig. S3. FACS gating strategy to select yeast cells that express RBD mutants with reduced binding by serum antibodies.

Fig. S4. Site- and mutation-level correlations between serum-escape measurements for each replicate library. Fig. S5. Binding-escape maps for the day 119 sera from all 14 individuals who received the 250 μg vaccine dose.

Fig. S6. Escape maps from individuals who received the 100 μg dose of mRNA-1273 119 days post-vaccination largely resemble those of individuals who received the 250 μg dose.

Fig. S7. Complete escape maps six representative convalescent plasmas from the day 100–150 cohort.

Fig. S8. Escape maps and effects of individual RBD mutations on neutralization for representative samples from vaccinated and convalescent individuals.

Fig. S9. Full neutralization curves for all assays testing how RBD mutations affected viral neutralization. Table S1. Serum neutralization titers pre- and post-depletion of RBD-binding antibodies.

Table S2. Information on FACS sorting to select cells expressing RBD mutants with reduced binding by sera from vaccinated individuals.

Table S3. Measurements of effects of all amino-acid mutations to the RBD on serum binding.

## Acknowledgments

We thank Cathy Lin for administrative support; Dolores Covarrubias, Andy Marty, and the Genomics and Flow Cytometry core facilities at the Fred Hutchinson Cancer Research Center for experimental support. This work used samples from the phase 1 mRNA-1273 study (NCT04283461; DOI: 10.1056/NEJMoa2022483). The mRNA-1273 phase 1 study was sponsored and primarily funded by the National Institute of Allergy and Infectious Diseases (NIAID), National Institutes of Health (NIH), Bethesda, MD; this trial has been funded in part with federal funds from the NIAID under grant awards UM1AI148373, to Kaiser Washington; UM1AI148576, UM1AI148684, and NIH P51 OD011132, to Emory University; NIH AID AI149644, and contract award HHSN272201500002C, to Emmes. Funding for the manufacture of mRNA-1273 phase 1 material was provided by the Coalition for Epidemic Preparedness Innovation. We also thank all research participants and study staff of the Hospitalized or Ambulatory Adults with Respiratory Viral Infections (HAARVI) study.

## Funding

National Institutes of Health R01AI141707 (JDB) National Institutes of Health R01AI127893 (JDB) National Institutes of Health T32AI083203 (AJG) National Institutes of Health F30AI149928 (KHDC) Gates Foundation INV-004949 (JDB)

National Institutes of Health Office of Research Infrastructure Programs S10OD028685 (The Scientific Computing Infrastructure at Fred Hutch).

Howard Hughes Medical Institute Fellow of the Damon Runyon Cancer Research Foundation DRG-2381-19 (TNS)

Howard Hughes Medical Institute (JDB) Gates Foundation INV-016575 (HYC) Emergent Ventures Award (HYC)

The content is solely the responsibility of the authors and does not necessarily represent the official views of the US government or the other sponsors.

## Author contributions

Conceptualization: AJG, JDB Methodology: AJG, TNS, KHDC Investigation: AJG, ANL, LEG, KDM Visualization: AJG, JDB

Resources: HYC, LAJ

Funding acquisition: JDB, HYC Supervision: JDB, HYC

Writing – original draft: AJG, JDB Writing – review & editing: all authors

## Data and materials availability

### Resource Availability

Further information and requests for reagents and resources should be directed to and will be fulfilled by Jesse Bloom.

#### Materials Availability

The SARS-CoV-2 RBD mutant libraries (#1000000172) and unmutated parental plasmid (#166782) are available on Addgene. The plasmid encoding the SARS-CoV-2 spike gene used to generate pseudotyped lentiviral particles, HDM_Spikedelta21_D614G, is available from Addgene (#158762) and BEI (NR-53765).

#### Data and Code Availability

We provide data and code in the following ways:

- The complete code for the full computational data analysis pipeline of the mapping experiments is available on GitHub at https://github.com/jbloomlab/SARS-CoV-2-RBD_MAP_Moderna.
- The escape fraction measured for each mutation in **Supplementary Table 3** and also at https://github.com/jbloomlab/SARS-CoV-2-RBD_MAP_Moderna/blob/main/results/supp_data/Moderna_convalescent_all_raw_data.csv.
- All raw sequencing data are available on the NCBI Short Read Archive at BioProject PRJNA639956, BioSample SAMN18683769.
- The neutralization titers of vaccine- and infection-elicited sera against the tested RBD point mutants is at https://github.com/jbloomlab/SARS-CoV-2-RBD_MAP_Moderna/blob/main/experimental_data/results/mutant_neuts_results/fitparams.csv.

---

**Fig. S1.**
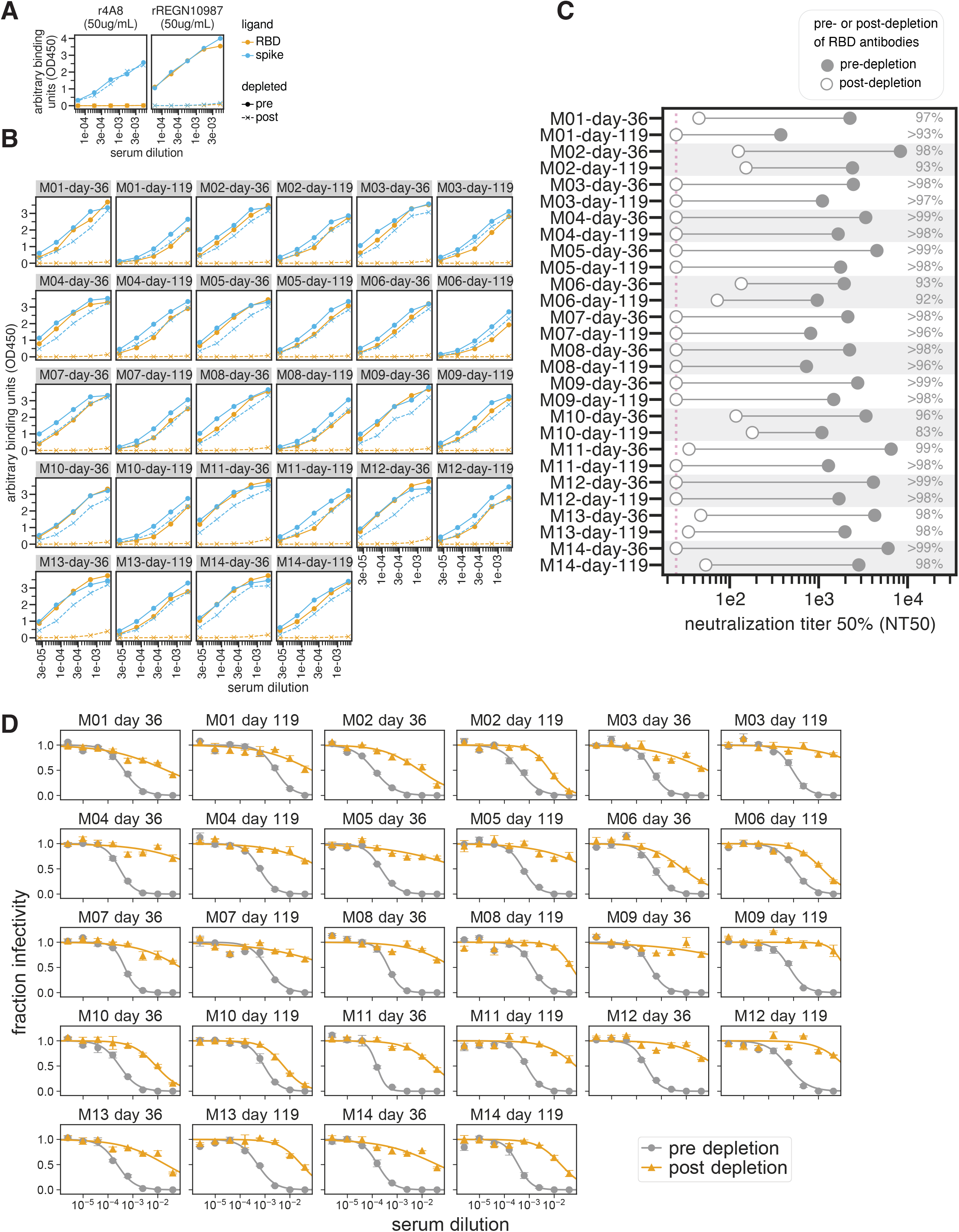
Raw ELISA and neutralization curves of mRNA-1273 serum samples before and after depletion of RBD-binding antibodies. **(A)** Effect of RBD antibody depletion on binding to RBD and spike by “synthetic sera'’ comprised of pre-pandemic pooled serum with the NTD-targeting antibody r4A8 (*21*) or RBD-targeting antibody rREGN10987 (*49*). Antibodies were added to pre-pandemic serum at 50 μg/mL. The x-axis indicates the dilution factor of the serum+antibody mix, and the y-axis is the ELISA reading at each dilution. These controls were previously used in (*15*), and demonstrate that the depletions effectively remove RBD-targeting antibodies but not antibodies targeting other epitopes such as the NTD. **(B)** Raw ELISA binding curves of sera to RBD and spike before and after depletion of RBD-binding antibodies. Legend for panels **(A)** and **(B)**: orange is RBD binding, blue is spike binding; filled circles with solid lines represent pre-depletion, and x’s with dashed lines represent post-depletion of anti-RBD antibodies. **(C)** Neutralization titer 50% (NT50) of vaccine-elicited sera pre- and post-depletion of RBD-binding antibodies, shown in filled and open circles, respectively. All neutralization assays were performed with SARS-CoV-2 spike D614G-pseudotyped lentiviral particles. Two time points were assessed per individual, at day 36 and day 119 post-dose 1 of vaccination. The limit of detection is shown as a dashed pink vertical line. The percent neutralization due to RBD-binding antibodies are shown at right. **(D)** Raw neutralization curves for sera before (gray) and after (orange) depletion of RBD-binding antibodies. Each assay was performed in technical duplicate, and points show the mean and standard error of the replicates.

**Fig. S2.**
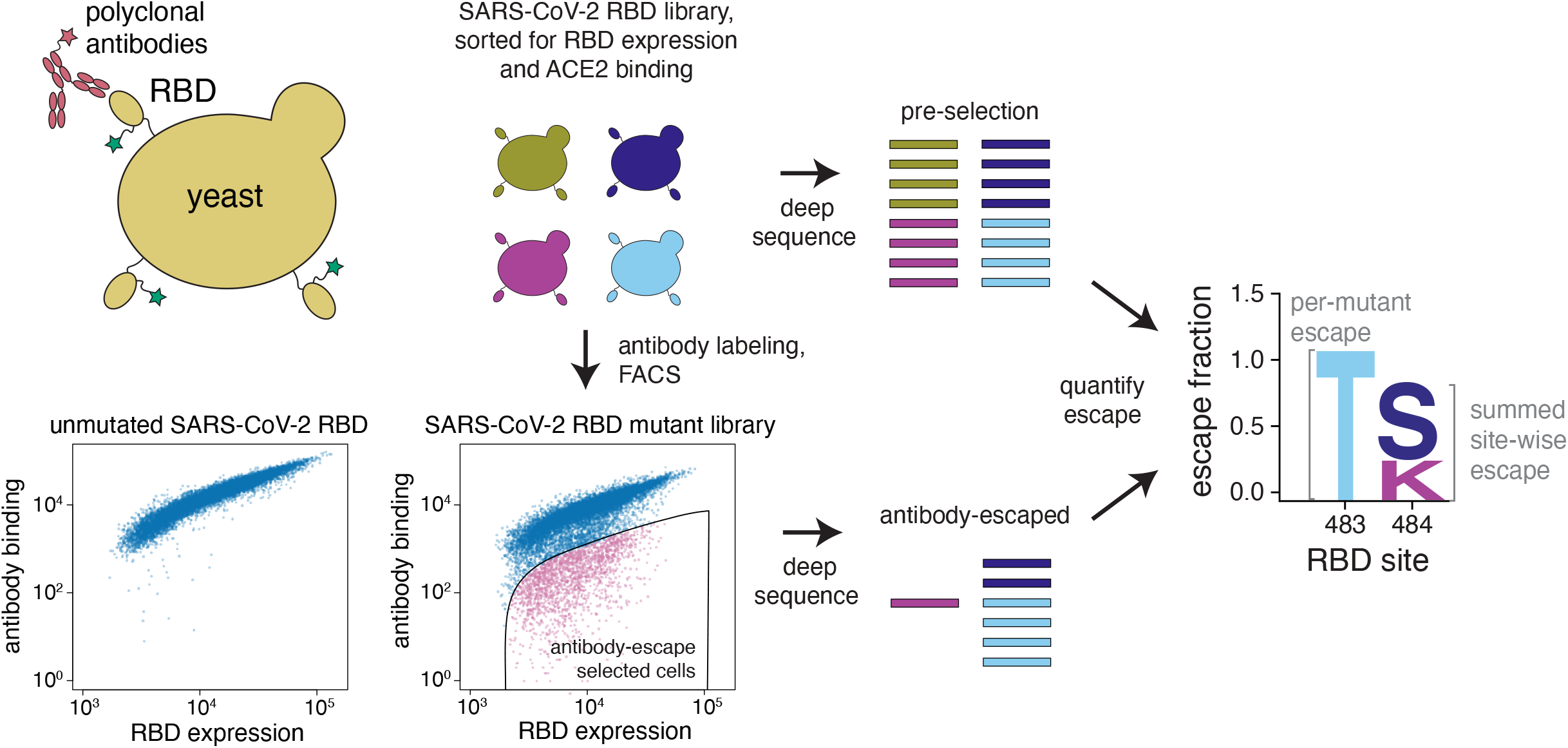
Schematic of the deep mutational scanning approach used to quantify the effects of RBD mutations on antibody escape. The RBD is expressed on the surface of yeast (top left). Flow cytometry is used to quantify both RBD expression (via a C-terminal MYC tag) and antibody binding to the RBD protein expressed on the surface of each yeast cell (bottom left). A library of yeast expressing RBD mutants was incubated with polyclonal serum and fluorescence-activated cell sorting (FACS) was used to enrich for cells expressing RBD that bound reduced levels of serum antibodies, as detected using an IgA+IgG+IgM secondary antibody. Deep sequencing was used to quantify the frequency of each mutation in the initial and “antibody escape” cell populations. We quantified the effect of each mutation as the “escape fraction,” which represents the fraction of cells expressing RBD with that mutation that fell in the “antibody escape” FACS bin. Escape fractions are represented in logo plots, with the height of each letter proportional to the effect of that amino-acid mutation on antibody binding. The site-level escape metric is the sum of the escape fractions of all mutations at a site. Experimental and computational filtering was used to remove RBD mutants that were misfolded or unable to bind the ACE2 receptor (see **Methods**).

**Fig. S3.**
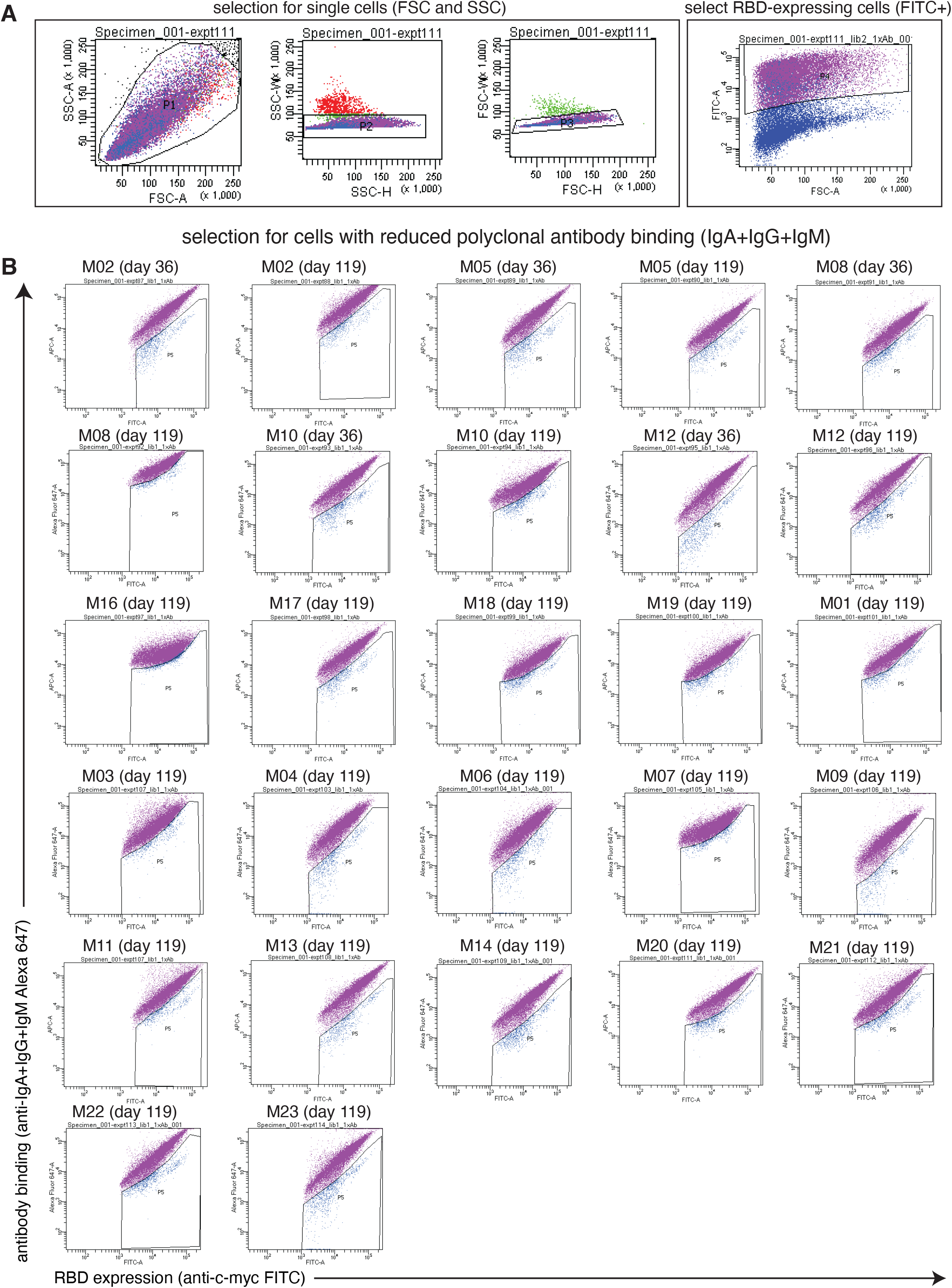
FACS gating strategy to select yeast cells that express RBD mutants with reduced binding by serum antibodies. **(A)** Representative plots of nested FACS gating strategy used for all serum selection experiments to select for single cells (SSC-A vs. FSC-A, SSC-W vs. SSC-H, and FSC-W vs. FSC-H) that also express RBD (FITC-A vs. FSC-A). **(B)** FACS gating strategy for one of two independent libraries to select cells expressing RBD mutants with reduced binding by polyclonal sera (cells in blue). Gates were set manually during sorting. Selection gates were set to capture ~5% of the RBD+ library. The same gate was set for both independent libraries stained with each serum, and the FACS scatter plots looked qualitatively similar between the two libraries. For information on the fraction of library cells that fall into each selection gate, see **Supplementary Table 2.**

**Fig. S4.**
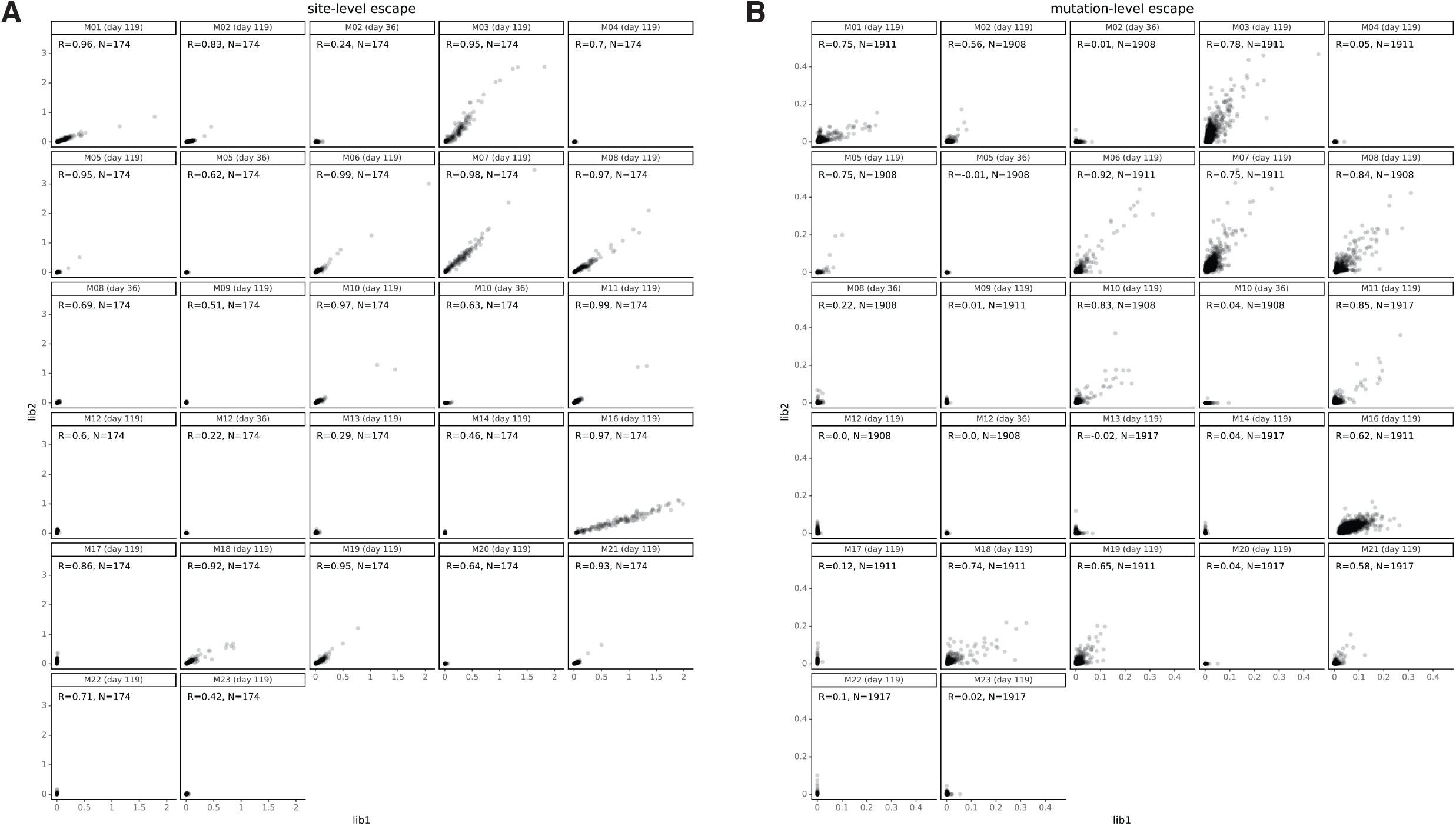
Site- and mutation-level correlations between serum-escape measurements for each replicate library. Correlation plots of site- or mutation-level escape for each of the two independent RBD mutant libraries for each serum sample. Each point represents one site in the RBD in **(A)** or a different mutation in **(B)**.

**Fig. S5.**
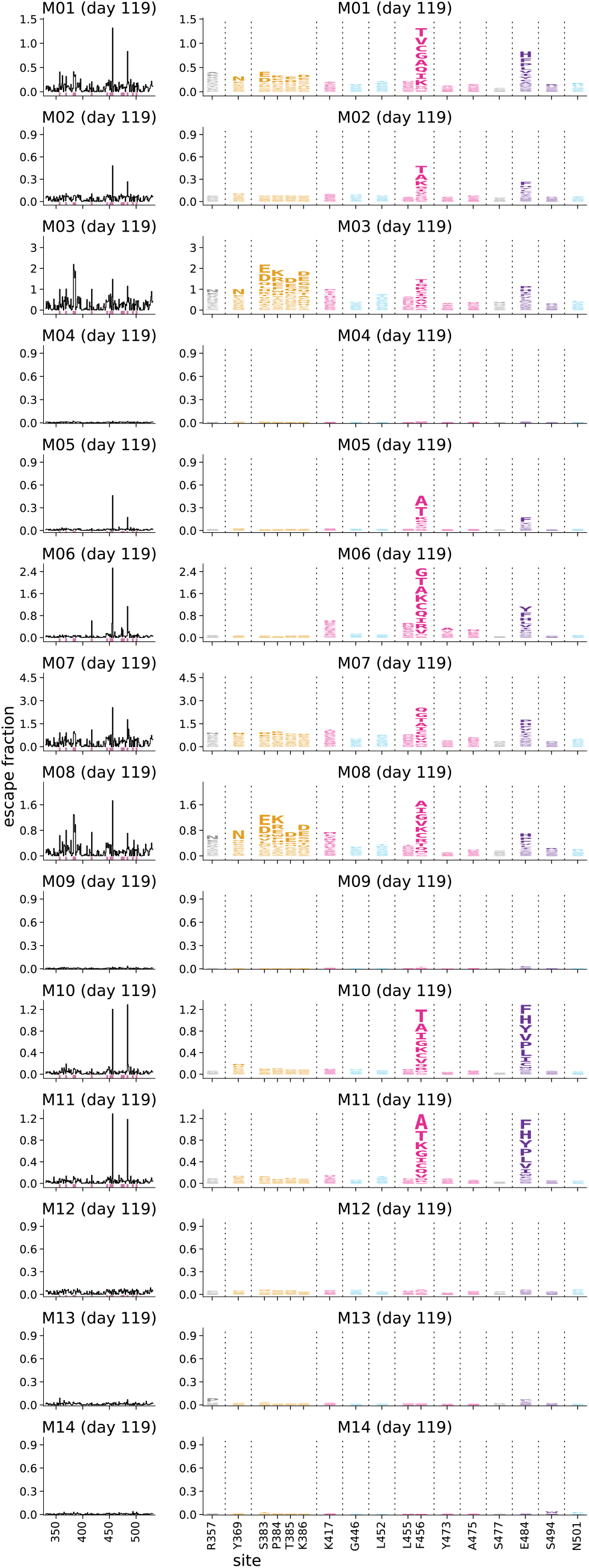
Binding-escape maps for the day 119 sera from all 14 individuals who received the 250 μg vaccine dose. Complete escape maps for the day 119 sera from the 14 individuals who received the 250 μg dose of mRNA-1273; note that a subset of these sera are also shown in **Fig. 2C**. RBD sites are colored according to epitope, as in **Fig. 2A**.

**Fig.S6.**
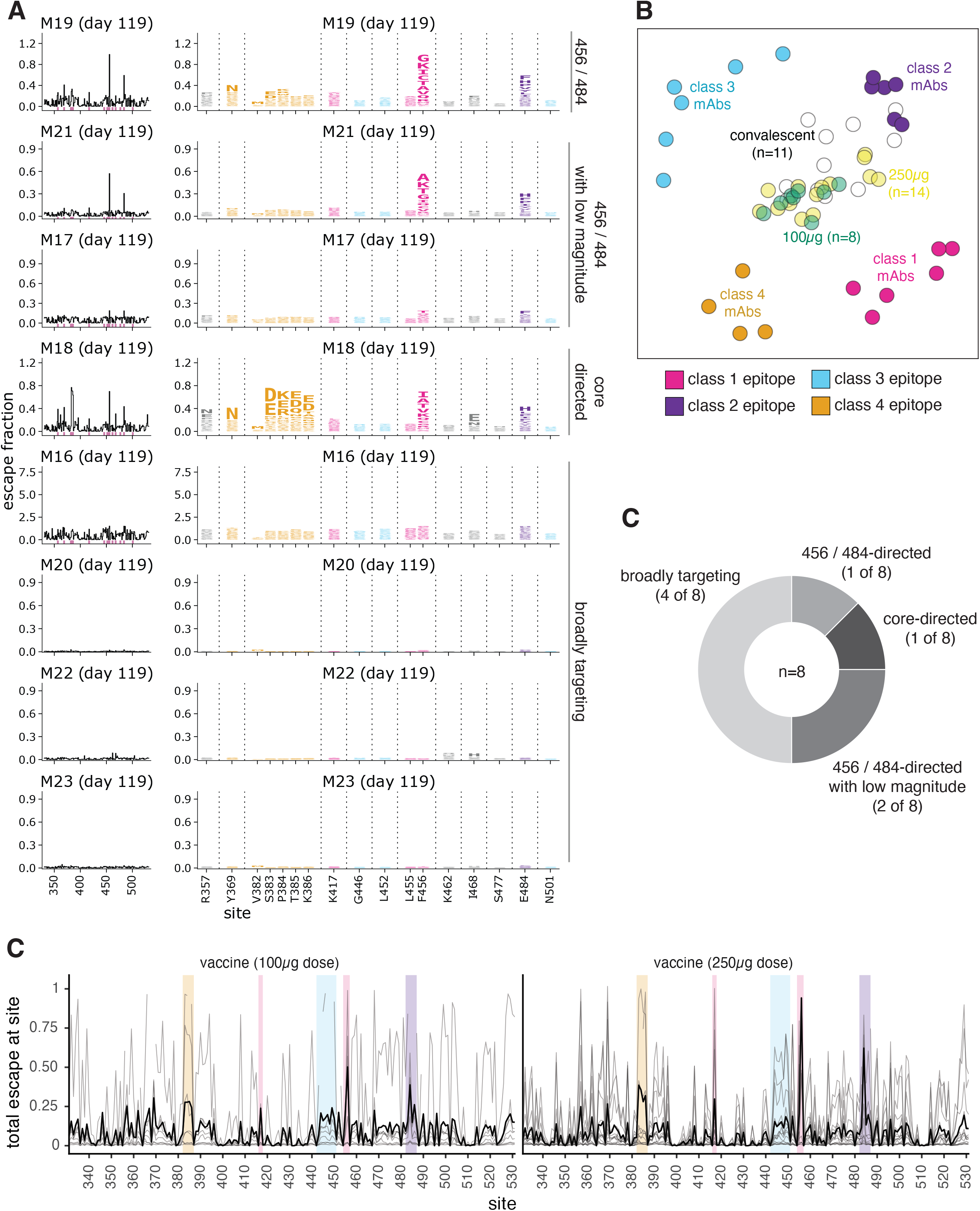
Escape maps from individuals who received the 100 μg dose of mRNA-1273 119 days post-vaccination largely resemble those of individuals who received the 250 μg dose. **(A)** Logo plots representing the complete escape maps for the day 119 sera from 8 individuals who received the 100 μg dose of mRNA-1273. RBD sites are colored according to epitope, as in **Fig. 2A**. **(B)** Multidimensional scaling projection that illustrates relationships among escape maps of sera and monoclonal antibodies in two dimensions, similarly to the projection shown in **Fig. 4**. Similar mutations affect the binding of antibodies or sera located near one another in the plot. Here, the only serum samples shown are the day 119 samples for individuals who received the 100 or 250 μg vaccine dose. The 100 μg dose samples are shown in green and the 250 μg dose samples are shown in yellow (day 119 for all). The projection includes the escape maps of 22 monoclonal antibodies (escape maps first described in (*16*, *22*, *25*–*27*) of the 4 major structural classes to orient the plot. Antibodies are colored according to epitope, as in **Fig. 2A**. **(C)** Composite line plots showing the total binding escape at each site in the RBD for each sample in gray, and the mean site-total escape for each group (100 or 250 μg vaccine doses) as a thicker black line. 100 μg, n=8; 250 μg, n=14. The same key sites within each epitope are highlighted, as in **Fig. 4**.

**Fig. S7.**
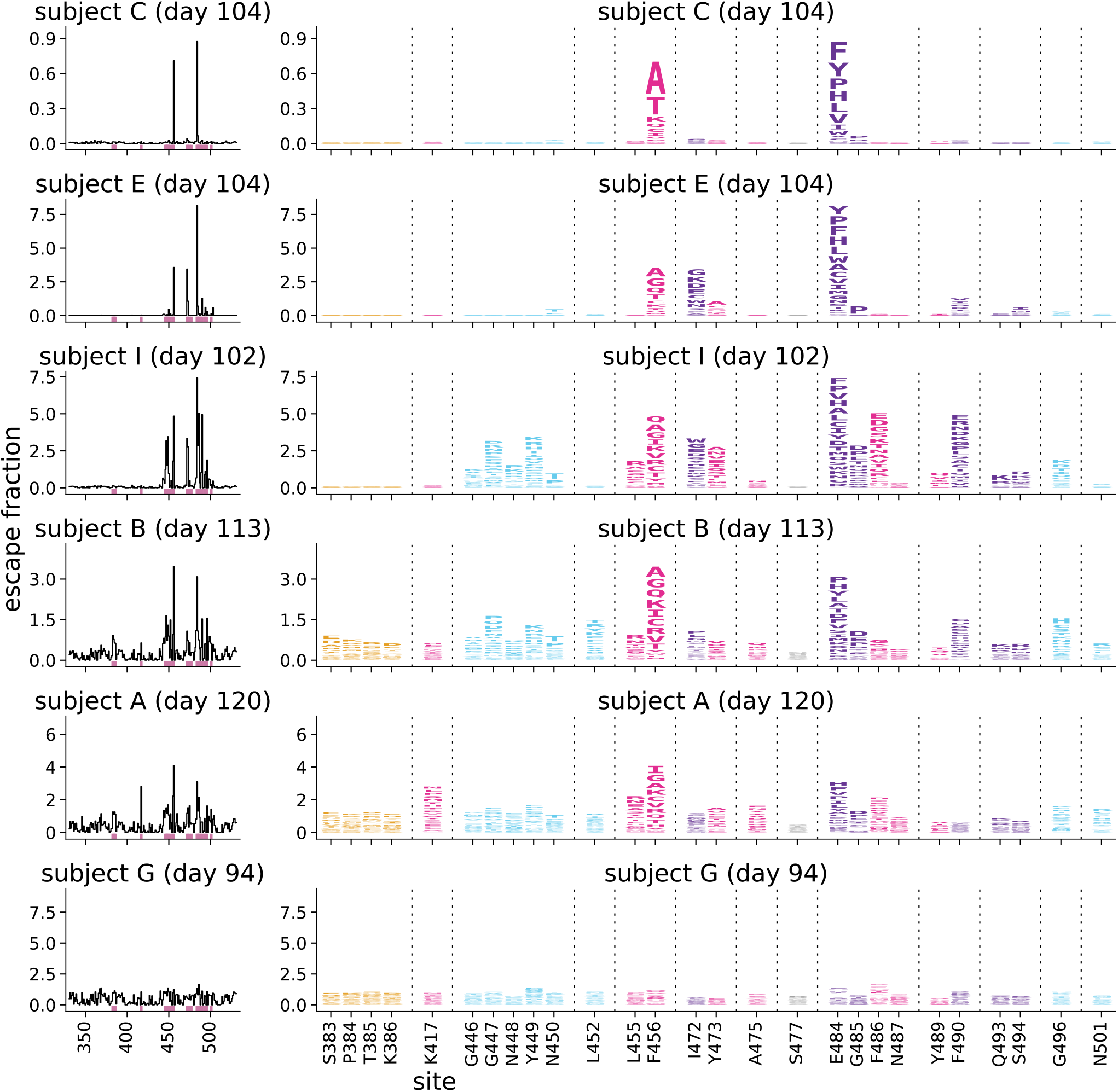
Complete escape maps six representative convalescent plasmas from the day 100–150 cohort. RBD sites are colored according to epitope, as in **Fig. 2A**. The same sites are shown here as in **Fig. 2A**. These six plasmas are those used in neutralization assays shown in **Fig. 5, S 8, S 9**. Escape maps were first reported in (*15*).

**Fig. S8.**
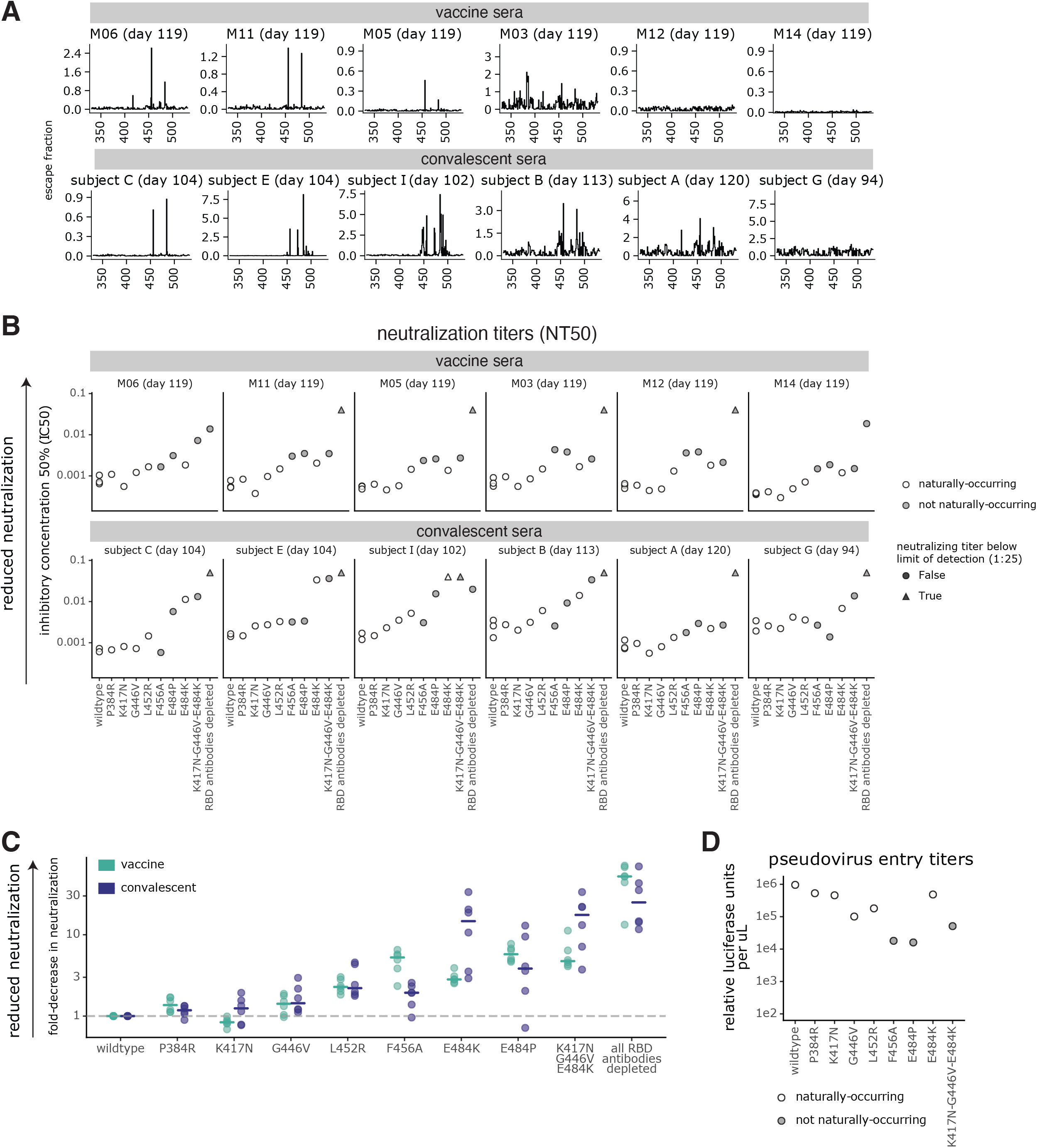
Escape maps and effects of individual RBD mutations on neutralization for representative samples from vaccinated and convalescent individuals. **(A)** The site-wise antibody-binding escape for each of the vaccine and convalescent samples tested in neutralization assays in panels B and C and Fig. 5. **(B)** The effects of RBD mutations on neutralization of G614 spike-pseudotyped lentiviral particles with the indicated mutations, shown as the inhibitory concentration 50% (IC50). Naturally-occurring mutations are colored in white, and non-naturally-occurring mutations in gray. **(C)** The fold-change in IC50 compared to wild type spike, grouped by vaccine or convalescent sera (as in Fig. 5C, but shown here for all RBD mutants). Dashed line indicates no change in neutralization relative to wild type spike. Horizontal bars represent the group median fold-change IC50. In (B) and (C), each point represents the IC50 from one individual calculated from technical duplicates. The highest two points for E484K and K417N-G446V-E484K, and the highest 4 points for “all RBD antibodies depleted” are at the limit of detection. **(D)** Spike-pseudotyped lentiviral particle entry titers for RBD mutants tested in neutralization assays, calculated as the mean relative luciferase units per μL from 16 technical replicates. Mutations that are observed in at least one SARS-CoV-2 sequence in GISAID are colored in white, and non-naturally-occurring mutations in gray. All spike sequences contained G614, which fixed in circulating sequences in 2020 (*47*). All full neutralization curves are in **Fig. S9** and raw IC50 and NT50 values are at https://github.com/jbloomlab/SARS-CoV-2-RBD_MAP_Moderna/blob/main/experimental_data/results/mutant_neuts_results/fitparams.csv.

**Fig. S9.**
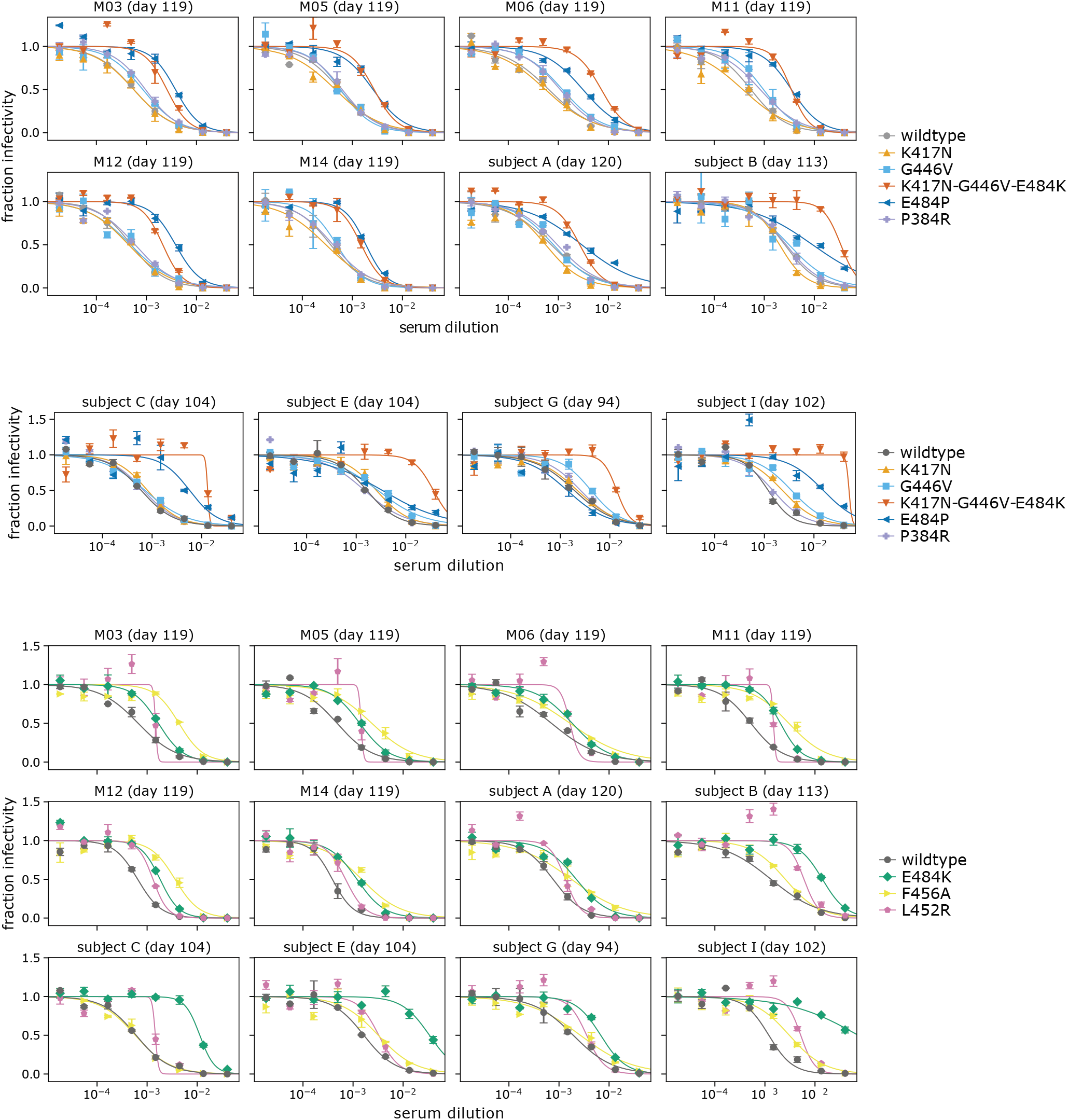
Full neutralization curves for all assays testing how RBD mutations affected viral neutralization. The x-axis is the serum dilution, and the y-axis is the fraction of viral infectivity remaining at that dilution. The neutralization curves were fit and plotted using neutcurve (https://jbloomlab.github.io/neutcurve/, version 0.5.2). IC50s were calculated by fitting 2-parameter Hill curves with the baselines fixed at one and zero. These IC50s were used to determine the fold-change values in **Fig. 5** and **S8**. In each plot, mutants are shown with the wildtype tested on the same date. Error bars represent the standard error of n=2 replicates. For readability, no more than 6 curves are shown per plot. In **Fig. 5D**, the wildtype curve from the first assay date is shown. Neutralization titers are available at https://github.com/jbloomlab/SARS-CoV-2-RBD_MAP_Moderna/blob/main/experimental_data/results/mutant_neuts_results/fitparams.csv

## Supplementary Files

**Table S1. Serum neutralization titers pre- and post-depletion of RBD-binding antibodies**. This file contains the serum neutralization titers from vaccinated individuals before (NT50_pre) and after (NT50_post) depletion of RBD-binding antibodies. The table is available online at https://github.com/jbloomlab/SARS-CoV-2-RBD_MAP_Moderna/blob/main/experimental_data/results/rbd_absorptions/TableS1.csv.

**Table S2. Information on FACS sorting to select cells expressing RBD mutants with reduced binding by sera from vaccinated individuals.** The file gives the number of antibody-escaped cells collected per selection for each replicate library and the percent of RBD+ cells in the antibody-escape gate for each selection, and the exact dilution used for each serum selection. The file is also available on GitHub at https://github.com/jbloomlab/SARS-CoV-2-RBD_MAP_Moderna/blob/main/data/TableS2_FACSinfo.csv.

**Table S3. Measurements of effects of all amino-acid mutations to the RBD on serum binding.**

The file gives the “escape fraction” for each mutation, as well as the total escape fraction at each site and the maximum escape fraction for any mutation at the site. This file includes escape fractions for sera from individuals vaccinated with mRNA-1273 as well as the previously reported escape fractions for convalescent plasma (*15*, *16*). The file is also available on GitHub at https://github.com/jbloomlab/SARS-CoV-2-RBD_MAP_Moderna/blob/main/results/supp_data/moderna_convalescent_all_raw_data.csv.

## References and Notes

1. O. W. Schmidt, I. D. Allan, M. K. Cooney, H. M. Foy, J. P. Fox, Rises in titers of antibody to human coronaviruses OC43 and 229E in Seattle families during 1975-1979. Am. J. Epidemiol. 123, 862–868 (1986).

2. R. T. Eguia, K. H. D. Crawford, T. Stevens-Ayers, L. Kelnhofer-Millevolte, A. L. Greninger, J. A. Englund, M. J. Boeckh, J. D. Bloom, A human coronavirus evolves antigenically to escape antibody immunity. PLoS Pathog. 17, e1009453 (2021).

3. S. Cele, I. Gazy, L. Jackson, S.-H. Hwa, H. Tegally, G. Lustig, J. Giandhari, S. Pillay, E. Wilkinson, Y. Naidoo, F. Karim, Y. Ganga, K. Khan, M. Bernstein, A. B. Balazs, B. I. Gosnell, W. Hanekom, M.-Y. S. Moosa, NGS-SA, COMMIT-KZN Team, R. J. Lessells, T. de Oliveira, A. Sigal, Escape of SARS-CoV-2 501Y.V2 from neutralization by convalescent plasma. Nature, 2021.01.26.21250224 (2021).

4. W. F. Garcia-Beltran, E. C. Lam, K. St Denis, A. D. Nitido, Z. H. Garcia, B. M. Hauser, J. Feldman, M. N. Pavlovic, D. J. Gregory, M. C. Poznansky, A. Sigal, A. G. Schmidt, A. J. Iafrate, V. Naranbhai, A. B. Balazs, Multiple SARS-CoV-2 variants escape neutralization by vaccine-induced humoral immunity. Cell (2021), doi:10.1016/j.cell.2021.03.013.

5. Z. Wang, F. Schmidt, Y. Weisblum, F. Muecksch, C. O. Barnes, S. Finkin, D. Schaefer-Babajew, M. Cipolla, C. Gaebler, J. A. Lieberman, T. Y. Oliveira, Z. Yang, M. E. Abernathy, K. E. Huey-Tubman, A. Hurley, M. Turroja, K. A. West, K. Gordon, K. G. Millard, V. Ramos, J. D. Silva, J. Xu, R. A. Colbert, R. Patel, J. Dizon, C. Unson-O’Brien, I. Shimeliovich, A. Gazumyan, M. Caskey, P. J. Bjorkman, R. Casellas, T. Hatziioannou, P. D. Bieniasz, M. C. Nussenzweig, mRNA vaccine-elicited antibodies to SARS-CoV-2 and circulating variants. Nature, 2021.01.15.426911 (2021).

6. R. E. Chen, X. Zhang, J. B. Case, E. S. Winkler, Y. Liu, L. A. VanBlargan, J. Liu, J. M. Errico, X. Xie, N. Suryadevara, P. Gilchuk, S. J. Zost, S. Tahan, L. Droit, J. S. Turner, W. Kim, A. J. Schmitz, M. Thapa, D. Wang, A. C. M. Boon, R. M. Presti, J. A. O’Halloran, A. H. J. Kim, P. Deepak, D. Pinto, D. H. Fremont, J. E. Crowe Jr, D. Corti, H. W. Virgin, A. H. Ellebedy, P.-Y. Shi, M. S. Diamond, Resistance of SARS-CoV-2 variants to neutralization by monoclonal and serum-derived polyclonal antibodies. Nat. Med. (2021), doi:10.1038/s41591-021-01294-w.

7. P. Wang, M. S. Nair, L. Liu, S. Iketani, Y. Luo, Y. Guo, M. Wang, J. Yu, B. Zhang, P. D. Kwong, B. S. Graham, J. R. Mascola, J. Y. Chang, M. T. Yin, M. Sobieszczyk, C. A. Kyratsous, L. Shapiro, Z. Sheng, Y. Huang, D. D. Ho, Antibody Resistance of SARS-CoV-2 Variants B.1.351 and B.1.1.7. Nature (2021), doi:10.1038/s41586-021-03398-2.

8. C. K. Wibmer, F. Ayres, T. Hermanus, M. Madzivhandila, P. Kgagudi, B. Oosthuysen, B. E. Lambson, T. de Oliveira, M. Vermeulen, K. van der Berg, T. Rossouw, M. Boswell, V. Ueckermann, S. Meiring, A. von Gottberg, C. Cohen, L. Morris, J. N. Bhiman, P. L. Moore, SARS-CoV-2 501Y.V2 escapes neutralization by South African COVID-19 donor plasma. Nat. Med., 2021.01.18.427166 (2021).

9. Novavax COVID-19 Vaccine Demonstrates 89.3% Efficacy in UK Phase 3 Trial, (available at https://ir.novavax.com/news-releases/news-release-details/novavax-covid-19-vaccine-demonstrates-893-efficacy-uk-phase-3).

10. Pfizer and BioNTech Confirm High Efficacy and No Serious Safety Concerns Through Up to Six Months Following Second Dose in Updated Topline Analysis of Landmark COVID-19 Vaccine Study, (available at https://www.pfizer.com/news/press-release/press-release-detail/pfizer-and-biontech-confirm-high-efficacy-and-no-serious).

11. K. S. Corbett, D. K. Edwards, S. R. Leist, O. M. Abiona, S. Boyoglu-Barnum, R. A. Gillespie, S. Himansu, A. Schäfer, C. T. Ziwawo, A. T. DiPiazza, K. H. Dinnon, S. M. Elbashir, C. A. Shaw, A. Woods, E. J. Fritch, D. R. Martinez, K. W. Bock, M. Minai, B. M. Nagata, G. B. Hutchinson, K. Wu, C. Henry, K. Bahl, D. Garcia-Dominguez, L. Ma, I. Renzi, W.-P. Kong, S. D. Schmidt, L. Wang, Y. Zhang, E. Phung, L. A. Chang, R. J. Loomis, N. E. Altaras, E. Narayanan, M. Metkar, V. Presnyak, C. Liu, M. K. Louder, W. Shi, K. Leung, E. S. Yang, A. West, K. L. Gully, L. J. Stevens, N. Wang, D. Wrapp, N. A. Doria-Rose, G. Stewart-Jones, H. Bennett, G. S. Alvarado, M. C. Nason, T. J. Ruckwardt, J. S. McLellan, M. R. Denison, J. D. Chappell, I. N. Moore, K. M. Morabito, J. R. Mascola, R. S. Baric, A. Carfi, B. S. Graham, SARS-CoV-2 mRNA vaccine design enabled by prototype pathogen preparedness. Nature. 586, 567–571 (2020).

12. L. A. Jackson, E. J. Anderson, N. G. Rouphael, P. C. Roberts, M. Makhene, R. N. Coler, M. P. McCullough, J. D. Chappell, M. R. Denison, L. J. Stevens, A. J. Pruijssers, A. McDermott, B. Flach, N. A. Doria-Rose, K. S. Corbett, K. M. Morabito, S. O’Dell, S. D. Schmidt, P. A. Swanson 2nd, M. Padilla, J. R. Mascola, K. M. Neuzil, H. Bennett, W. Sun, E. Peters, M. Makowski, J. Albert, K. Cross, W. Buchanan, R. Pikaart-Tautges, J. E. Ledgerwood, B. S. Graham, J. H. Beigel, mRNA-1273 Study Group, An mRNA Vaccine against SARS-CoV-2 - Preliminary Report. N. Engl. J. Med. 383, 1920–1931 (2020).

13. K. H. D. Crawford, A. S. Dingens, R. Eguia, C. R. Wolf, N. Wilcox, J. K. Logue, K. Shuey, A. M. Casto, B. Fiala, S. Wrenn, D. Pettie, N. P. King, A. L. Greninger, H. Y. Chu, J. D. Bloom, Dynamics of neutralizing antibody titers in the months after SARS-CoV-2 infection. J. Infect. Dis. (2020), doi:10.1093/infdis/jiaa618.

14. D. F. Robbiani, C. Gaebler, F. Muecksch, J. C. C. Lorenzi, Z. Wang, A. Cho, M. Agudelo, C. O. Barnes, A. Gazumyan, S. Finkin, T. Hägglöf, T. Y. Oliveira, C. Viant, A. Hurley, H.-H. Hoffmann, K. G. Millard, R. G. Kost, M. Cipolla, K. Gordon, F. Bianchini, S. T. Chen, V. Ramos, R. Patel, J. Dizon, I. Shimeliovich, P. Mendoza, H. Hartweger, L. Nogueira, M. Pack, J. Horowitz, F. Schmidt, Y. Weisblum, E. Michailidis, A. W. Ashbrook, E. Waltari, J. E. Pak, K. E. Huey-Tubman, N. Koranda, P. R. Hoffman, A. P. West Jr, C. M. Rice, T. Hatziioannou, P. J. Bjorkman, P. D. Bieniasz, M. Caskey, M. C. Nussenzweig, Convergent antibody responses to SARS-CoV-2 in convalescent individuals. Nature. 584, 437–442 (2020).

15. A. J. Greaney, A. N. Loes, K. H. D. Crawford, T. N. Starr, K. D. Malone, H. Y. Chu, J. D. Bloom, Comprehensive mapping of mutations in the SARS-CoV-2 receptor-binding domain that affect recognition by polyclonal human plasma antibodies. Cell Host Microbe (2021), doi:10.1016/j.chom.2021.02.003.

16. A. J. Greaney, T. N. Starr, C. O. Barnes, Y. Weisblum, F. Schmidt, M. Caskey, C. Gaebler, M. Agudelo, S. Finkin, Z. Wang, D. Poston, F. Muecksch, T. Hatziioannou, P. D. Bieniasz, D. F. Robbiani, M. C. Nussenzweig, P. J. Bjorkman, J. D. Bloom, Mutational escape from the polyclonal antibody response to SARS-CoV-2 infection is largely shaped by a single class of antibodies. bioRxiv, 2021.03.17.435863 (2021).

17. L. Piccoli, Y.-J. Park, M. A. Tortorici, N. Czudnochowski, A. C. Walls, M. Beltramello, C. Silacci-Fregni, D. Pinto, L. E. Rosen, J. E. Bowen, O. J. Acton, S. Jaconi, B. Guarino, A. Minola, F. Zatta, N. Sprugasci, J. Bassi, A. Peter, A. De Marco, J. C. Nix, F. Mele, S. Jovic, B. F. Rodriguez, S. V. Gupta, F. Jin, G. Piumatti, G. Lo Presti, A. F. Pellanda, M. Biggiogero, M. Tarkowski, M. S. Pizzuto, E. Cameroni, C. Havenar-Daughton, M. Smithey, D. Hong, V. Lepori, E. Albanese, A. Ceschi, E. Bernasconi, L. Elzi, P. Ferrari, C. Garzoni, A. Riva, G. Snell, F. Sallusto, K. Fink, H. W. Virgin, A. Lanzavecchia, D. Corti, D. Veesler, Mapping Neutralizing and Immunodominant Sites on the SARS-CoV-2 Spike Receptor-Binding Domain by Structure-Guided High-Resolution Serology. Cell. 183, 1024–1042.e21 (2020).

18. W. Dejnirattisai, D. Zhou, H. M. Ginn, H. M. E. Duyvesteyn, P. Supasa, J. B. Case, Y. Zhao, T. S. Walter, A. J. Mentzer, C. Liu, B. Wang, G. C. Paesen, J. Slon-Campos, C. López-Camacho, N. M. Kafai, A. L. Bailey, R. E. Chen, B. Ying, C. Thompson, J. Bolton, A. Fyfe, S. Gupta, T. K. Tan, J. Gilbert-Jaramillo, W. James, M. Knight, M. W. Carroll, D. Skelly, C. Dold, Y. Peng, R. Levin, T. Dong, A. J. Pollard, J. C. Knight, P. Klenerman, N. Temperton, D. R. Hall, M. A. Williams, N. G. Paterson, F. K. R. Bertram, C. A. Seibert, D. K. Clare, A. Howe, J. Raedecke, Y. Song, A. R. Townsend, K.-Y. A. Huang, E. E. Fry, J. Mongkolsapaya, M. S. Diamond, J. Ren, D. I. Stuart, G. R. Screaton, The antigenic anatomy of SARS-CoV-2 receptor binding domain. Cell (2021), doi:10.1016/j.cell.2021.02.032.

19. F. Amanat, M. Thapa, T. Lei, S. M. S. Ahmed, D. C. Adelsberg, J. M. Carreno, S. Strohmeier, A. J. Schmitz, S. Zafar, J. Q. Zhou, W. Rijnink, H. Alshammary, N. Borcherding, A. G. Reiche, K. Srivastava, E. M. Sordillo, H. van Bakel, J. S. Turner, G. Bajic, V. Simon, A. H. Ellebedy, F. Krammer, The Personalized Virology Initiative, The plasmablast response to SARS-CoV-2 mRNA vaccination is dominated by non-neutralizing antibodies that target both the NTD and the RBD. bioRxiv (2021), , doi:10.1101/2021.03.07.21253098.

20. N. Suryadevara, S. Shrihari, P. Gilchuk, L. A. VanBlargan, E. Binshtein, S. J. Zost, R. S. Nargi, R. E. Sutton, E. S. Winkler, E. C. Chen, M. E. Fouch, E. Davidson, B. J. Doranz, R. E. Chen, P.-Y. Shi, R. H. Carnahan, L. B. Thackray, M. S. Diamond, J. E. Crowe Jr, Neutralizing and protective human monoclonal antibodies recognizing the N-terminal domain of the SARS-CoV-2 spike protein. Cell, 2021.01.19.427324 (2021).

21. X. Chi, R. Yan, J. Zhang, G. Zhang, Y. Zhang, M. Hao, Z. Zhang, P. Fan, Y. Dong, Y. Yang, Z. Chen, Y. Guo, J. Zhang, Y. Li, X. Song, Y. Chen, L. Xia, L. Fu, L. Hou, J. Xu, C. Yu, J. Li, Q. Zhou, W. Chen, A neutralizing human antibody binds to the N-terminal domain of the Spike protein of SARS-CoV-2. Science. 369, 650–655 (2020).

22. A. J. Greaney, T. N. Starr, P. Gilchuk, S. J. Zost, E. Binshtein, A. N. Loes, S. K. Hilton, J. Huddleston, R. Eguia, K. H. D. Crawford, A. S. Dingens, R. S. Nargi, R. E. Sutton, N. Suryadevara, P. W. Rothlauf, Z. Liu, S. P. J. Whelan, R. H. Carnahan, J. E. Crowe Jr, J. D. Bloom, Complete Mapping of Mutations to the SARS-CoV-2 Spike Receptor-Binding Domain that Escape Antibody Recognition. Cell Host Microbe. 29, 44–57.e9 (2021).

23. T. N. Starr, A. J. Greaney, S. K. Hilton, D. Ellis, K. H. D. Crawford, A. S. Dingens, M. J. Navarro, J. E. Bowen, M. A. Tortorici, A. C. Walls, N. P. King, D. Veesler, J. D. Bloom, Deep Mutational Scanning of SARS-CoV-2 Receptor Binding Domain Reveals Constraints on Folding and ACE2 Binding. Cell. 182, 1295–1310.e20 (2020).

24. C. O. Barnes, C. A. Jette, M. E. Abernathy, K.-M. A. Dam, S. R. Esswein, H. B. Gristick, A. G. Malyutin, N. G. Sharaf, K. E. Huey-Tubman, Y. E. Lee, D. F. Robbiani, M. C. Nussenzweig, A. P. West Jr, P. J. Bjorkman, SARS-CoV-2 neutralizing antibody structures inform therapeutic strategies. Nature. 588, 682–687 (2020).

25. T. N. Starr, A. J. Greaney, A. Addetia, W. W. Hannon, M. C. Choudhary, A. S. Dingens, J. Z. Li, J. D. Bloom, Prospective mapping of viral mutations that escape antibodies used to treat COVID-19. Science (2021), doi:10.1126/science.abf9302.

26. T. N. Starr, A. J. Greaney, A. S. Dingens, J. D. Bloom, Complete map of SARS-CoV-2 RBD mutations that escape the monoclonal antibody LY-CoV555 and its cocktail with LY-CoV016. Cell Reports Medicine, 100255 (2021).

27. J. Dong, S. J. Zost, A. J. Greaney, T. N. Starr, A. S. Dingens, E. C. Chen, R. E. Chen, J. B. Case, R. E. Sutton, P. Gilchuk, J. Rodriguez, E. Armstrong, C. Gainza, R. S. Nargi, E. Binshtein, X. Xie, X. Zhan, P.-Y. Shi, J. Logue, S. Weston, M. E. McGrath, M. B. Frieman, T. Brady, K. Tuffy, H. Bright, Y.-M. Loo, P. McTamney, M. Esser, R. H. Carnahan, M. S. Diamond, J. D. Bloom, J. E. Crowe, Genetic and structural basis for recognition of SARS-CoV-2 spike protein by a two-antibody cocktail. bioRxiv, 2021.01.27.428529 (2021).

28. N. R. Faria, I. M. Claro, D. Candido, L. A. Moyses Franco, P. S. Andrade, T. M. Coletti, C. A. M. Silva, F. C. Sales, E. R. Manuli, R. S. Aguiar, N. Gaburo, C. da C. Camilo, N. A. Fraiji, M. A. Esashika Crispim, M. do Perpétuo S. S. Carvalho, A. Rambaut, N. Loman, O. G. Pybus, E. C. Sabino, on behalf of CADDE Genomic Network, Genomic characterisation of an emergent SARS-CoV-2 lineage in Manaus: preliminary findings. Virological (2021) (available at https://virological.org/t/genomic-characterisation-of-an-emergent-sars-cov-2-lineage-in-manaus-preliminary-findings/586).

29. C. M. Voloch, R. da Silva Francisco Jr, L. G. P. de Almeida, C. C. Cardoso, O. J. Brustolini, A. L. Gerber, A. P. de C. Guimarães, D. Mariani, R. M. da Costa, O. C. Ferreira Jr, Covid19-UFRJ Workgroup, LNCC Workgroup, Adriana Cony Cavalcanti, T. S. Frauches, C. M. B. de Mello, I. de C. Leitão, R. M. Galliez, D. S. Faffe, T. M. P. P. Castiñeiras, A. Tanuri, A. T. R. de Vasconcelos, Genomic characterization of a novel SARS-CoV-2 lineage from Rio de Janeiro, Brazil. J. Virol. (2021), doi:10.1128/JVI.00119-21.

30. W. Zhang, B. D. Davis, S. S. Chen, J. M. Sincuir Martinez, J. T. Plummer, E. Vail, Emergence of a Novel SARS-CoV-2 Variant in Southern California. JAMA (2021), doi:10.1001/jama.2021.1612.

31. A. P. West, C. O. Barnes, Z. Yang, P. J. Bjorkman, SARS-CoV-2 lineage B.1.526 emerging in the New York region detected by software utility created to query the spike mutational landscape. bioRxiv, 2021.02.14.431043 (2021).

32. M. K. Annavajhala, H. Mohri, J. E. Zucker, Z. Sheng, P. Wang, A. Gomez-Simmonds, D. D. Ho, A.-C. Uhlemann, A novel SARS-CoV-2 variant of concern, B.1.526, identified in New York. medRxiv (2021), , doi:10.1101/2021.02.23.21252259.

33. H. Tegally, E. Wilkinson, M. Giovanetti, A. Iranzadeh, V. Fonseca, J. Giandhari, D. Doolabh, S. Pillay, E. J. San, N. Msomi, K. Mlisana, A. von Gottberg, S. Walaza, M. Allam, A. Ismail, T. Mohale, A. J. Glass, S. Engelbrecht, G. Van Zyl, W. Preiser, F. Petruccione, A. Sigal, D. Hardie, G. Marais, M. Hsiao, S. Korsman, M.-A. Davies, L. Tyers, I. Mudau, D. York, C. Maslo, D. Goedhals, S. Abrahams, O. Laguda-Akingba, A. Alisoltani-Dehkordi, A. Godzik, C. K. Wibmer, B. T. Sewell, J. Lourenço, L. C. J. Alcantara, S. L. Kosakovsky Pond, S. Weaver, D. Martin, R. J. Lessells, J. N. Bhiman, C. Williamson, T. de Oliveira, Emergence of a SARS-CoV-2 variant of concern with mutations in spike glycoprotein. Nature (2021), doi:10.1038/s41586-021-03402-9.

34. D. Planas, T. Bruel, L. Grzelak, F. Guivel-Benhassine, I. Staropoli, F. Porrot, C. Planchais, J. Buchrieser, M. M. Rajah, E. Bishop, M. Albert, F. Donati, S. Behillil, V. Enouf, M. Maquart, M. Gonzalez, J. De Sèze, H. Péré, D. Veyer, A. Sève, E. Simon-Lorière, S. Fafi-Kremer, K. Stefic, H. Mouquet, L. Hocqueloux, S. van der Werf, T. Prazuck, O. Schwartz, Sensitivity of infectious SARS-CoV-2 B.1.1.7 and B.1.351 variants to neutralizing antibodies. bioRxiv, 2021.02.12.430472 (2021).

35. M. McCallum, J. Bassi, A. De Marco, A. Chen, A. C. Walls, J. Di Iulio, M. Alejandra Tortorici, M.-J. Navarro, C. Silacci-Fregni, C. Saliba, M. Agostini, D. Pinto, K. Culap, S. Bianchi, S. Jaconi, E. Cameroni, J. E. Bowen, S. W. Tiles, M. S. Pizzuto, S. B. Guastalla, G. Bona, A. F. Pellanda, C. Garzoni, W. C. Van Voorhis, L. E. Rosen, G. C. Snell, A. Telenti, H. W. Virgin, L. Piccoli, D. Corti, D. Veesler, SARS-CoV-2 immune evasion by variant B.1.427/B.1.429. bioRxiv (2021), p. 2021.03.31.437925.

36. S. J. Zost, P. Gilchuk, J. B. Case, E. Binshtein, R. E. Chen, J. P. Nkolola, A. Schäfer, J. X. Reidy, A. Trivette, R. S. Nargi, R. E. Sutton, N. Suryadevara, D. R. Martinez, L. E. Williamson, E. C. Chen, T. Jones, S. Day, L. Myers, A. O. Hassan, N. M. Kafai, E. S. Winkler, J. M. Fox, S. Shrihari, B. K. Mueller, J. Meiler, A. Chandrashekar, N. B. Mercado, J. J. Steinhardt, K. Ren, Y.-M. Loo, N. L. Kallewaard, B. T. McCune, S. P. Keeler, M. J. Holtzman, D. H. Barouch, L. E. Gralinski, R. S. Baric, L. B. Thackray, M. S. Diamond, R. H. Carnahan, J. E. Crowe Jr, Potently neutralizing and protective human antibodies against SARS-CoV-2. Nature. 584, 443–449 (2020).

37. L. Liu, P. Wang, M. S. Nair, J. Yu, M. Rapp, Q. Wang, Y. Luo, J. F.-W. Chan, V. Sahi, A. Figueroa, X. V. Guo, G. Cerutti, J. Bimela, J. Gorman, T. Zhou, Z. Chen, K.-Y. Yuen, P. D. Kwong, J. G. Sodroski, M. T. Yin, Z. Sheng, Y. Huang, L. Shapiro, D. D. Ho, Potent neutralizing antibodies against multiple epitopes on SARS-CoV-2 spike. Nature. 584, 450–456 (2020).

38. A. Kuzmina, Y. Khalaila, O. Voloshin, A. Keren-Naus, L. Bohehm, Y. Raviv, Y. Shemer-Avni, E. Rosenberg, R. Taube, SARS CoV-2 escape variants exhibit differential infectivity and neutralization sensitivity to convalescent or post-vaccination sera. bioRxiv (2021), doi:10.1101/2021.02.22.21252002.

39. C. Gaebler, Z. Wang, J. C. C. Lorenzi, F. Muecksch, S. Finkin, M. Tokuyama, M. Ladinsky, A. Cho, M. Jankovic, D. Schaefer-Babajew, T. Y. Oliveira, M. Cipolla, C. Viant, C. O. Barnes, A. Hurley, M. Turroja, K. Gordon, K. G. Millard, V. Ramos, F. Schmidt, Y. Weisblum, D. Jha, M. Tankelevich, J. Yee, I. Shimeliovich, D. F. Robbiani, Z. Zhao, A. Gazumyan, T. Hatziioannou, P. J. Bjorkman, S. Mehandru, P. D. Bieniasz, M. Caskey, M. C. Nussenzweig, Evolution of Antibody Immunity to SARS-CoV-2. Nature (2020), doi:10.1038/s41586-021-03207-w.

40. F. Muecksch, Y. Weisblum, C. Barnes, F. Schmidt, D. Schaefer-Babajew, J. Lorenzi, A. Flyak, A. DeLaitsch, K. Huey-Tubman, S. Hou, C. Schiffer, C. Gaebler, Z. Wang, J. Da Silva, D. Poston, S. Finkin, A. Cho, M. Cipolla, T. Oliveira, K. Millard, V. Ramos, A. Gazumyan, M. Rutkowska, M. Caskey, M. Nussenzweig, P. Bjorkman, T. Hatziioannou, P. Bieniasz, Development of potency, breadth and resilience to viral escape mutations in SARS-CoV-2 neutralizing antibodies. bioRxiv, 2021.03.07.434227 (2021).

41. L. Dai, G. F. Gao, Viral targets for vaccines against COVID-19. Nat. Rev. Immunol. 21, 73–82 (2021).

42. N. Pardi, M. J. Hogan, F. W. Porter, D. Weissman, mRNA vaccines - a new era in vaccinology. Nat. Rev. Drug Discov. 17, 261–279 (2018).

43. K. Roeltgen, S. C. A. Nielsen, P. S. Arunachalam, F. Yang, R. A. Hoh, O. F. Wirz, A. S. Lee, F. Gao, V. Mallajosyula, C. Li, E. Haraguchi, M. J. Shoura, J. L. Wilbur, J. N. Wohlstadter, M. M. Davis, B. A. Pinsky, G. B. Sigal, B. Pulendran, K. C. Nadeau, S. D. Boyd, mRNA vaccination compared to infection elicits an IgG-predominant response with greater SARS-CoV-2 specificity and similar decrease in variant spike recognition. medRxiv, 2021.04.05.21254952 (2021).

44. A. J. Greaney, A. N. Loes, K. H. D. Crawford, T. N. Starr, K. D. Malone, H. Y. Chu, J. D. Bloom, Comprehensive mapping of mutations to the SARS-CoV-2 receptor-binding domain that affect recognition by polyclonal human serum antibodies. bioRxiv (2021), p. 2020.12.31.425021.

45. J. Otwinowski, D. M. McCandlish, J. B. Plotkin, Inferring the shape of global epistasis. Proc. Natl. Acad. Sci. U. S. A. 115, E7550–E7558 (2018).

46. K. H. D. Crawford, R. Eguia, A. S. Dingens, A. N. Loes, K. D. Malone, C. R. Wolf, H. Y. Chu, M. A. Tortorici, D. Veesler, M. Murphy, D. Pettie, N. P. King, A. B. Balazs, J. D. Bloom, Protocol and Reagents for Pseudotyping Lentiviral Particles with SARS-CoV-2 Spike Protein for Neutralization Assays. Viruses. 12 (2020), doi:10.3390/v12050513.

47. B. Korber, W. M. Fischer, S. Gnanakaran, H. Yoon, J. Theiler, W. Abfalterer, N. Hengartner, E. E. Giorgi, T. Bhattacharya, B. Foley, K. M. Hastie, M. D. Parker, D. G. Partridge, C. M. Evans, T. M. Freeman, T. I. de Silva, Sheffield COVID-19 Genomics Group, C. McDanal, L. G. Perez, H. Tang, A. Moon-Walker, S. P. Whelan, C. C. LaBranche, E. O. Saphire, D. C. Montefiori, Tracking Changes in SARS-CoV-2 Spike: Evidence that D614G Increases Infectivity of the COVID-19 Virus. Cell. 182, 812–827.e19 (2020).

48. A. S. Dingens, K. H. D. Crawford, A. Adler, S. L. Steele, K. Lacombe, R. Eguia, F. Amanat, A. C. Walls, C. R. Wolf, M. Murphy, D. Pettie, L. Carter, X. Qin, N. P. King, D. Veesler, F. Krammer, J. A. Dickerson, H. Y. Chu, J. A. Englund, J. D. Bloom, Serological identification of SARS-CoV-2 infections among children visiting a hospital during the initial Seattle outbreak. Nat. Commun. 11, 4378 (2020).

49. J. Hansen, A. Baum, K. E. Pascal, V. Russo, S. Giordano, E. Wloga, B. O. Fulton, Y. Yan, K. Koon, K. Patel, K. M. Chung, A. Hermann, E. Ullman, J. Cruz, A. Rafique, T. Huang, J. Fairhurst, C. Libertiny, M. Malbec, W.-Y. Lee, R. Welsh, G. Farr, S. Pennington, D. Deshpande, J. Cheng, A. Watty, P. Bouffard, R. Babb, N. Levenkova, C. Chen, B. Zhang, A. Romero Hernandez, K. Saotome, Y. Zhou, M. Franklin, S. Sivapalasingam, D. C. Lye, S. Weston, J. Logue, R. Haupt, M. Frieman, G. Chen, W. Olson, A. J. Murphy, N. Stahl, G. D. Yancopoulos, C. A. Kyratsous, Studies in humanized mice and convalescent humans yield a SARS-CoV-2 antibody cocktail. Science. 369, 1010–1014 (2020).

50. J. Lan, J. Ge, J. Yu, S. Shan, H. Zhou, S. Fan, Q. Zhang, X. Shi, Q. Wang, L. Zhang, X. Wang, Structure of the SARS-CoV-2 spike receptor-binding domain bound to the ACE2 receptor. Nature. 581, 215–220 (2020).

51. S. Hilton, J. Huddleston, A. Black, K. North, A. Dingens, T. Bedford, J. Bloom, dms-view: Interactive visualization tool for deep mutational scanning data. J. Open Source Softw. 5, 2353 (2020).

